# Constructed wetlands for aquaculture wastewater treatment: insights on the structural and functional shifts of the aquatic microbial community

**DOI:** 10.1101/2025.09.02.673640

**Authors:** D. Corso, M. Melita, N. Massaccesi, G. M. Quero, M. Basili, A. Di Cesare, R. Sabatino, T. Sbaffi, S. Fazi, A. Rakaj, G. M. Luna, S. Amalfitano

## Abstract

Aquaculture practices generate nutrient-rich effluents with associated microbiological hazards, such as pathogens and antimicrobial resistance genes (ARGs). Despite their growing popularity as nature-based solutions, little is known about how constructed wetlands (CWs) affect the dynamics of microbial communities at the field scale. By combining flow cytometry, 16S rRNA gene sequencing, shotgun metagenomics, and metabolic potential assays, we investigated the structural and functional responses of the aquatic microbial community following the recurrent exposure to CW-treated effluents from an intensive marine fish farm (Orbetello lagoon, Italy). While the CW promoted abundant, metabolically active, and functionally redundant microbial communities, the phylogenetic composition diverged primarily between water and sediments. Microbial profiles in CW outlet waters converged towards those of the lagoon baselines, suggesting gradual ecological recovery. The CW attenuated the occurrence of potential pathogens (e.g., *Francisella* spp., *Campylobacter* spp.) and limited ARG dissemination, though sediments remained reservoirs of microbial and genetic signatures. Functional profiles, dominated by chemoheterotrophy, denitrification, and sulfur respiration, remained stable across environments, reflecting microbial resilience. Our results highlight CWs as effective, field-proven solutions to mitigate aquaculture wastewater impacts while preserving core ecosystem services.

## 1. Introduction

Aquaculture has rapidly expanded in response to the increasing global demand for fish, driven by the human population growth and the limitation of wild fish catch production (Ahmad et al., 2022). This industrial sector is also facing environmental challenges, especially regarding the chemical and microbiological pollution of surrounding ecosystems. Aquaculture wastewater is rich in nutrients, organic matter, feed waste, and metabolic byproducts, which can directly contribute to the eutrophication processes of receiving water bodies (Liu et al., 2024). Among the various strategies available to mitigate environmental and public health risks posed by nutrient-enriched aquaculture effluents, constructed wetlands (CWs) have been recognized as sustainable and cost-effective treatment systems (Kushwaha et al., 2024; Liu et al., 2024; Moazzem et al., 2023). CWs are nature-based solutions that replicate and promote natural remediation processes through a combination of physical, chemical, and biological mechanisms, including sedimentation, adsorption, biodegradation, and denitrification processes (Kushwaha et al., 2024; Moazzem et al., 2023). Particularly, the aquatic microbial communities play a crucial role in the biodegradation, removal, and transformation of pollutants and excess nutrients, also supported by local vegetation (e.g., phytodepuration), available molecules promoting redox reactions (e.g., oxygen, nitrate, sulfate), and biofilm formation at the water-sediment interface (Hassan et al., 2021; Moazzem et al., 2023).

The microbial assemblages in CWs are diverse, dynamic, and shaped by a range of factors, including abiotic parameters (e.g., temperature, pH, salinity, composition of wastewater, oxygenation level, seasonal variation), the operational settings of the CW system (e.g., lagooning period, hydraulic retention time, flow regime), and the biological properties of the influent water (Liu et al., 2024; Vymazal et al., 2021). Microbial diversity was proved crucial for effective pollutant removal, as different microbial groups are responsible for degrading specific contaminants through distinct metabolic pathways (Hassan et al., 2021). However, long-term pollution exposure can lead to a decrease in microbial diversity with the selection of pollutant-degrading microbial taxa, which may compromise the overall treatment performance (Xiong et al., 2023). The current understanding of microbial responses and community patterns is mostly grounded on CWs fed by urban runoff and municipal wastewater (Vymazal, 2022). Numerous studies emphasized the interconnections between system design, treatment performances, and their effects on microbial communities in both water and sediment (Verduzo Garibay et al., 2022a; Wang et al., 2022).

However, a significant gap remains in understanding how the aquatic microbial community shifts in response to recurrent exposure to aquaculture effluents (Verduzo Garibay et al., 2022b). The available studies were mainly conducted under controlled laboratory conditions using artificial wastewater and in small-scale outdoor systems (Vymazal et al., 2021). Field observations on the dynamics of microbial communities are likely essential for providing novel insights into the ecological impact of aquaculture practices and for improving the understanding of the effectiveness and sustainability of CWs in the treatment of intensive aquaculture effluents originating from full-scale productive systems (Ma et al., 2018; Niu et al., 2025; Tian et al., 2024).

The objective of this study was to explore the structural and functional responses of the aquatic microbial community after the treatment of aquaculture wastewater by a constructed wetland. More specifically, we aimed to (i) explore the microbial community patterns in water and sediment samples collected pre- and post-treatment, (ii) unveil the occurrence of potential pathogens and antimicrobial resistance genes, and (iii) assess the impact on key nutrient cycles (i.e., carbon, nitrogen, sulfur) in waters and sediments with long-term exposure to aquaculture effluents. Control samples collected from adjacent lagoon and coastal marine environments were used as ecological baselines. We hypothesize that CWs can promote the remediation of the aquaculture effluents by shifting the phylogenetic and functional profiles of the microbial community, as detected by high-throughput techniques such as flow cytometry, 16S rRNA gene amplicon sequencing, shotgun metagenomics, and metabolic potential assays.

## 2. Material and methods

### 2.1 Study area and sampling activities

This study targeted a constructed wetland of approximately 2.5 hectares, specifically designed to receive and treat the effluents of a land-based intensive fish farm located in the area of the Orbetello lagoon (Tuscany, Italy) (Figure 1). The aquaculture plant scheme is semi-closed and includes more than 60 fish tanks with an overall surface area of approx. 3.5 hectares, hosting adults and juveniles of sea bream (*Sparus aurata*), sea bass (*Dicentrarchus labrax*), and goldmouth croaker (*Argyrosomus regius*). The tanks with adult fish are approximately 2 meters deep and receive both saline groundwater and water recirculating through the CW pathway. Oxygen addition and major water quality parameters are monitored continuously by the fish farm managers to optimize the flow rate ratio between the two water sources while controlling ammonium and nutrient levels required for optimal fish growth.

**Figure 1.**
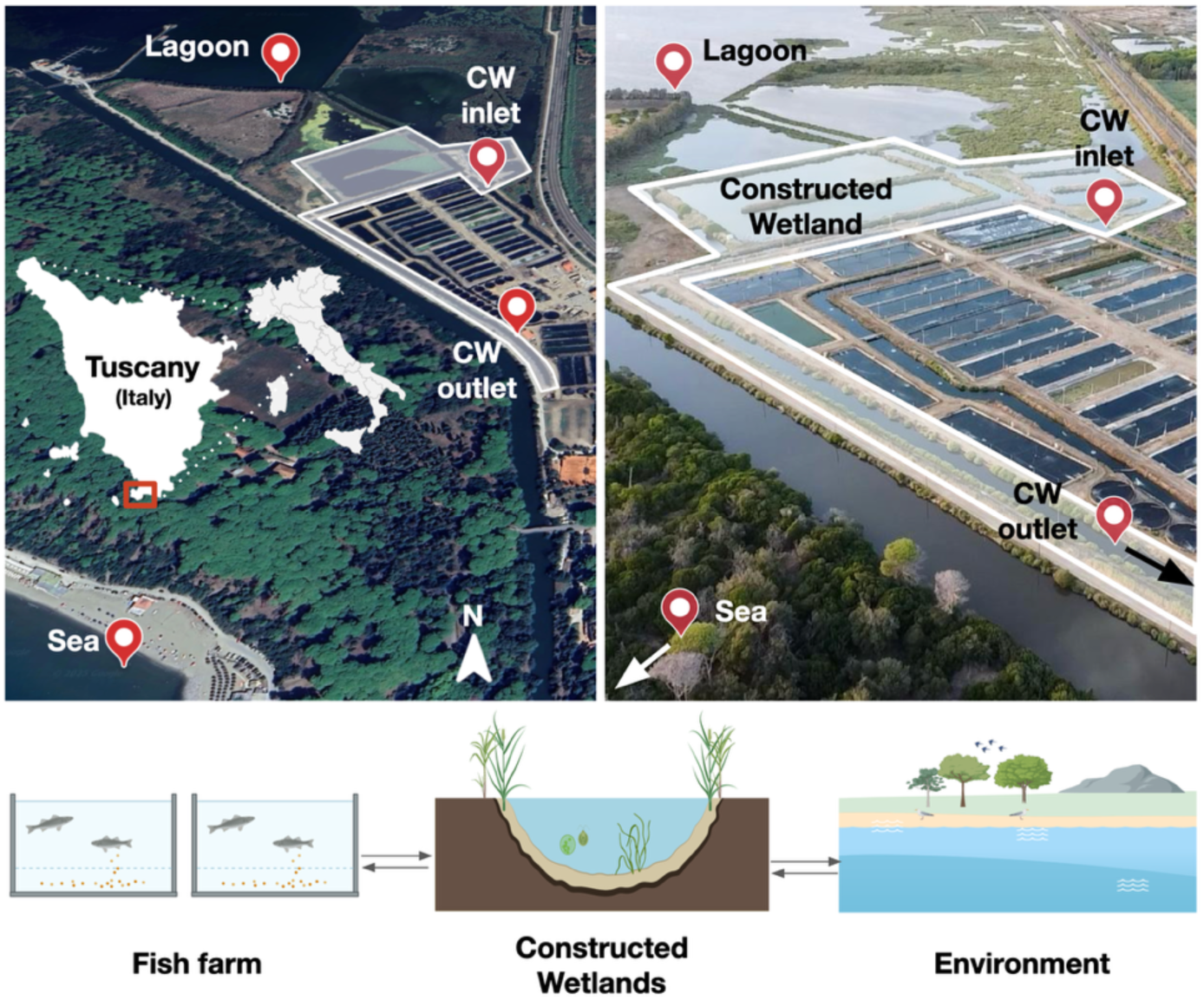
Constructed wetland (white shaded area) associated with a land-based intensive fish farm (Orbetello lagoon, Italy). Sampling points are marked in the satellite view (left panel). The CW area is highlighted in the aerial view of the aquaculture plant (right panel). Water and sediment samples were collected at the CW inlet (where aquaculture effluents enter the CW) and outlet (where CW-treated waters are discharged). Control samples were collected from the nearby lagoon and sea areas. The bottom shows the schematic relationships between the artificial and natural systems.

Sampling was carried out at the CW system, targeting both the inlet (i.e., where aquaculture effluents enter the CW) and the outlet (i.e., where CW-treated waters are discharged). To provide a comparative baseline, control samples were also collected from nearby non-aquaculture areas, including the adjacent coastal lagoon and the coastal marine environments, which were not directly affected by aquaculture effluents. Water samples were collected at each site using pre-cleaned 2-liter glass bottles. Surface sediment samples were collected using a hand-held telescopic scoop and transferred into clean plastic buckets, with approximately 1 kg of wet sediment obtained per sampling site. All samples were kept refrigerated and transported to the laboratory for subsequent processing on the same day of sampling. Field measurements were assessed three times from 11 AM to 6 PM during two sampling events in November 2023 (8-21/11/2023). A multiparameter field probe (HACH; HQ Series - Multi) was used at each sampling site to record basic physical-chemical parameters in water, including temperature (T), pH, electrical conductivity (EC), and dissolved oxygen (DO). Salinity (Sal.) was measured using a portable refractometer and expressed in parts per thousand (‰).

### 2.2. Microbial load assessment by flow cytometry

The abundance of prokaryotic cells and pigmented phytoplanktonic microbes (i.e., cyanobacteria, pico- and nano-eukaryotes) was assessed by the Flow Cytometer A50-micro (Apogee Flow System, Hertfordshire, UK), equipped with a 20 mW solid-state blue laser (488 nm). Water aliquots were immediately placed in 2-mL safe-lock tubes, fixed with formaldehyde (2% final concentration), and stored at 4°C. Sediment aliquots (1 g) were treated to detach cells in a liquid suspension following established procedures based on sonication/agitation in a buffered detergent solution (Amalfitano and Fazi, 2008). The light scattering signals (forward and side light scatter named FSC and SSC, respectively), red fluorescence (>610 nm), orange fluorescence (590/35 nm), and green fluorescence (535/35 nm) were acquired and considered for the direct identification and quantification of distinct microbial groups by following harmonized protocols (Amalfitano et al., 2018; Gasol and Morán, 2016). Cyanobacteria and pico-eukaryotic plankton were characterized and distinguished according to their pigmentation and size. Thresholding was set on the red channel to exclude most of the unspecific signals according to 0.2-μm filtered control water samples. The abundance of prokaryotic and nanoeukaryotic cells (e.g., bacteria and nanoflagellates) was determined by following the staining procedure with SYBR Green I (1:10,000 dilution; Molecular Probes, Life Technologies, code S7563) for 10 min in the dark at room temperature. Thresholding was set on the green channel, and the gating strategy was manually adjusted to exclude most of the unspecific signals according to negative unstained controls. Samples were run at low flow rates to keep the number of total events below 1,000 per second. Data handling and visualization were performed by the Apogee Histogram Software (v89.0).

### 2.3 Sample pre-treatment and DNA extraction

For DNA extraction, water samples (1 L) were filtered using 0.22 μm cellulose nitrate membranes (Sartorius, Ø 47 mm). The filter membranes were stored at −20°C until processing. Sediment samples were collected from each bucket using a sterile core extruder, from which 1 g was used for extraction. All extractions were performed using the DNeasy PowerSoil Kit (Qiagen), following the protocol described elsewhere (Quero et al., 2023). DNA concentration and purity were assessed using a NanoDrop ND-1000 spectrophotometer (NanoDrop Technologies, Wilmington, DE, USA), and samples were stored at −20°C prior to downstream analyses.

### 2.4 Phylogenetic community composition and functional annotation by 16S rRNA gene amplicon sequencing

The V3–V4 hypervariable region of the 16S rRNA gene was amplified using the primer pair 341F–785R (Klindworth et al., 2013), and the resulting PCR products were purified following the protocol described elsewhere (Basili et al., 2023). Library preparation was carried out using Nextera indexing, and sequencing was performed on the Illumina MiSeq platform with a 2 × 300 bp paired-end configuration as described in Basili et al. (2023).

The obtained reads were processed using the DADA2 pipeline for BigData paired-end reads (https://benjjneb.github.io/dada2/bigdata_paired.html) (Callahan et al., 2016) to retrieve amplicon sequence variants (ASVs). Figaro (Weinstein et al., 2019) and Cutadapt (Martin, 2011) were used to determine optimal trimming parameters and remove adapter or primer sequences from raw reads, following the tutorial at https://github.com/nuriamw/micro4all. The taxonomic annotation of the resulting ASVs was done using the SILVA reference database (version 138.2) (Quast et al., 2013). By integrating the taxonomic annotation and samples metadata, we analyzed the ASV abundance table through the R package phyloseq (McMurdie and Holmes, 2013). Raw counts data were normalized by median sequencing depth, and relative abundances were calculated. The data were then grouped at the family level and subsequently filtered to retain only major families, those with a relative abundance greater than 1% in at least one sample. To create a visual representation of the most relevant taxa, we kept classes that were found in at least three different samples. The sequencing data can be retrieved using the following BioProject ID on the NCBI database PRJNA1200612 (Supplementary Table 1).

The FAPROTAX database was used to annotate the functional metabolic potential of ASVs based on their taxonomy (Louca et al., 2016). Normalized counts by median sequencing depth were used as input. Next, we sorted the FAPROTAX results into groups based on possible metabolic functions related to the three main biogeochemical cycles (i.e., carbon, nitrogen, and sulfur). Moreover, we manually grouped the FAPROTAX functions associated with the occurrence of potential human, animal, and plant pathogens. Stacked bar plots were used to visualize the abundance of the major metabolic functions and the occurrence of potential pathogens at the genus level. We used online resources and three reference databases of medically relevant pathogenic bacteria to double-check the identified genera as potential pathogens (Virulence Factors Database - VFDB https://www.mgc.ac.cn/VFs/main.htm (Zhou et al., 2025), Bacterial and Viral Bioinformatics Resource Center - BV-BRC https://www.bv-brc.org (Olson et al., 2023), and BacDive https://bacdive.dsmz.de/about (Schober et al., 2025).

### 2.5 Antimicrobial resistance and key functional genes by shotgun metagenomics

Shotgun metagenomic sequencing was carried out by an external service (IGA Technology Services S.r.l., Udine, Italy) using indexed DNA libraries with an average fragment length of ∼500 bp. Libraries were sequenced on Illumina NextSeq instruments using NovaSeq 6000 reagents, producing 2 × 150 bp paired-end reads (300 cycles). The resulting raw reads were processed on the KBase platform (https://www.kbase.us/) (Arkin et al., 2018) following a comprehensive bioinformatic pipeline: (i) paired-end reads were merged; (ii) adapter sequences were removed using Trimmomatic (v0.39) (Bolger et al., 2014); (iii) quality control was performed with FastQC (v0.11.9) (Andrews, 2010); (iv) taxonomic classification of reads was carried out using Kaiju (Menzel et al., 2016); (v) antimicrobial resistance genes (ARGs) were identified on unassembled reads using CARD-RGI (Alcock et al., 2023), and both their occurrence and richness were assessed; (vi) reads were assembled with metaSPAdes (Nurk et al., 2017); (vii) read coverage was computed and assemblies were used for downstream analyses; (viii) ARGs on assembled contigs were again identified with CARD-RGI (Alcock et al., 2023); (ix) functional annotation of assembled contigs was performed using DRAM (Shaffer et al., 2023). The metagenomic sequencing data can be retrieved using the following BioProject ID on the NCBI database PRJNA1200612 (Supplementary Table 1).

### 2.6 Microbial metabolic potential and functional properties by BIOLOG assays

The Biolog™ EcoPlates assay (Biolog, Inc., Hayward, California, USA) was used to test the microbial metabolic potential and functional diversity in water and sediment samples, following 24 h incubation in 96 micro-well plates at 20°C (Melita et al., 2022). The colorimetric reduction of the respiration-sensitive dye tetrazolium to formazan upon microbial degradation was detected by measuring the absorbance at 590 nm through a spectrophotometric plate reader (PerkinElmer VICTOR™ X3 Multilabel Plate Reader). The assay was used to evaluate microbial mineralization of 31 carbon sources (tested in triplicate), grouped into six major classes of organic compounds: amines, amino acids, carbohydrates, carboxylic acids, phenols, and polymers, including 10 substrates containing aminic groups (–NH₂) (Supplementary Table 2). Microorganisms able to directly assimilate –NH₂ can incorporate organic nitrogen into their biomass through an energetically favorable process compared to nitrate reduction, since these molecules can be directly used for protein synthesis or deaminated to ammonium and released into the environment (Barros et al., 2023). The mean degradative activity across all 31 carbon substrates was used as a proxy of the microbial metabolic potential (Stefanowicz, 2006). The N-metabolic potential was assessed using only the 10 N-containing substrates. The functional diversity was estimated by calculating the Shannon Index from the degradation profiles of the different substrates (Miki et al., 2018).

### 2.7 Statistical analyses

Univariate and multivariate statistics were performed with the R programming language (version 4.5.0), using ggplot2 for the graphical visualization. Given the non-normal distribution of most data, the non-parametric Kruskal-Wallis test was employed to evaluate whether significant differences were observed for each microbial variable between water and sediment samples (waters vs sediments), as well as samples originating from the constructed wetland and the natural environmental controls (CW vs sea and lagoon). A significance threshold of p < 0.05 was applied for all tests. Alpha diversity with the Shannon diversity index was assessed using *plot_richness()* function applied to phylogenetic and metabolic data. Beta diversity was computed using non-parametric multidimensional scaling (NMDS) with Bray-Curtis and Gower as multiple distance metrics. The Bray–Curtis dissimilarity of phylogenetic data was calculated using either the ASVs or the family-level abundance table. A permutational multivariate analysis of variance (PERMANOVA with adonis2 function) was applied to investigate differences in the microbial community profiles between the two target groups (i.e., waters vs sediments; CW vs environment). Log-transformation [log(x+1)] was applied to normalize skewed data distributions. We used the *envfit* function on the same variables that were part of the NMDS input matrix, aiming to visualize the variables that were significantly related to the ordination structure (p < 0.05). The stress value was used to evaluate the overall quality of the NMDS ordination.

## 3. Results

### 3.1 Environmental parameters and microbial load

Field parameters showed consistent differences between waters from CW and those from the control environments (Table 1). The pH, conductivity, oxygen levels, and salinity remained consistent between the CW inlet and outlet. Higher values were found in the lagoon and sea waters. Over oxygenated waters were found in lagoon and CW outlet samples. Sediment composition was predominantly muddy at both the CW inlet and outlet, transitioning to a higher sand content in lagoon and coastal marine samples. One cubic centimeter of sediment was found to yield approximately two grams of wet material.

**Table 1.** Mean values (± standard deviation) of water parameters assessed during the sampling period (November 2023).

| Sample | Temp.<br>°C | pH | EC<br>mS/cm | DO<br>mg/L | DO<br>% | Sal.<br>‰ |
| --- | --- | --- | --- | --- | --- | --- |
| CW inlet | 20.8 ± 0.3 | 6.8 ± 0.1 | 51.7 ± 0.9 | 8.1 ± 0.1 | 90.8 ± 0.7 | 37.0 ± 0.0 |
| CW outlet | 20.1 ± 1.0 | 6.8 ± 0.0 | 51.8 ± 1.5 | 8.2 ± 2.2 | 90.8 ± 23.3 | 37.5 ± 0.5 |
| Lagoon | 18.2 ± 0.7 | 8.3 ± 0.1 | 53.0 ± 1.3 | 12.5 ± 0.5 | 132.2 ± 6.4 | 39.0 ± 1.0 |
| Sea | 20.0 ± 1.2 | 8.2 ± 0.0 | 55.0 ± 2.4 | 9.1 ± 0.3 | 99.0 ± 0.5 | 39.5 ± 0.5 |

Flow cytometry showed different microbial cell abundance and community structure in water and sediment between CW and the adjacent environments (Table 2).

**Table 2.**
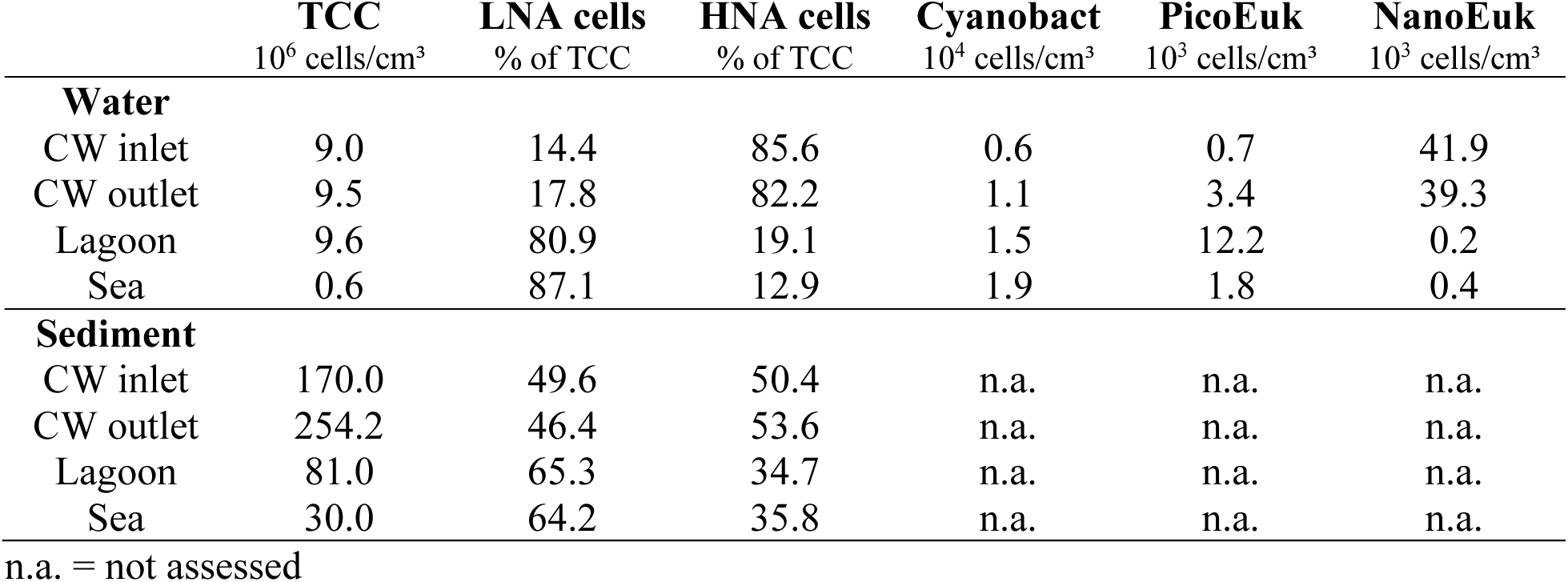
Microbial load in water and sediment samples. TCC = Total Cell Counts; LNA cells = cells with Low Nucleic Acid content; HNA cells = cells with High Nucleic Acid content; Cyanobact = Cyanobacteria; picoEuk = Picoeukaryotes (small eukaryotic plankton, 1–10 µm); NanoEuk = Nanoeukaryotes (larger eukaryotic plankton, 10–100 µm).

In water, TCC values were higher at the CW inlet, outlet, and lagoon compared to the sea. Sediments showed TCC values 10 to 100 times higher per unit of volume, especially at the CW outlet. LNA cells prevailed in the lagoon and sea, whereas HNA cells dominated at the CW inlet and outlet (> 80% of TCC). Sediment samples showed higher HNA percentages in CW sites but with a more balanced HNA/LNA cell ratio. Photosynthetic pigmented cells, mostly represented by cyanobacteria and picoeukaryotic plankton, increased from the CW to the lagoon and sea. Nanoeukaryotic plankton, consisting mostly of non-pigmented heterotrophic cells, was almost exclusively detected in CW waters.

### 3.2 Microbial community composition and functional annotation

By considering 46 phyla, 80 classes, 171 orders, and 339 families, distinct patterns of microbial community composition were found across sample types and locations, with marked differences between water and sediment samples, as well as between CW and control environments (lagoon and sea). The Kruskal-Wallis test applied to Shannon diversity indices showed non-significant results for both comparisons. PERMANOVA showed that the microbial community composition at the ASV level differed significantly between water and sediment samples (p = 0.031), but not between constructed wetlands and natural environments (p = 0.129). The dominant families (> 1% of total reads) belonged to 7 phyla, 9 classes, and 26 orders (Figure 2). The family Flavobacteriaceae (class Bacteroidia, phylum Bacteroidota) consistently exhibited high relative abundances across CW and environmental water samples, with abundance values ranging from 9.6% in seawater up to 29.3% in CW inlet water. Paracoccaceae (phylum Pseudomonadota, class Alphaproteobacteria) showed a high relative abundance in water samples, particularly in CW inlet (22.1%) and outlet (24.1%). In sediments, members of the phylum Thermodesulfobacteriota were the most abundant, with the families Desulfocapsaceae and Desulfosarcinaceae showing the highest relative abundance in the CW inlet and outlet. Other families were abundant only in specific environments. For example, the family Halieaceae (phylum Pseudomonadota, class Gammaproteobacteria) was abundant in lagoon sediments, reaching up to 22.8% of the total reads.

**Figure 2.**
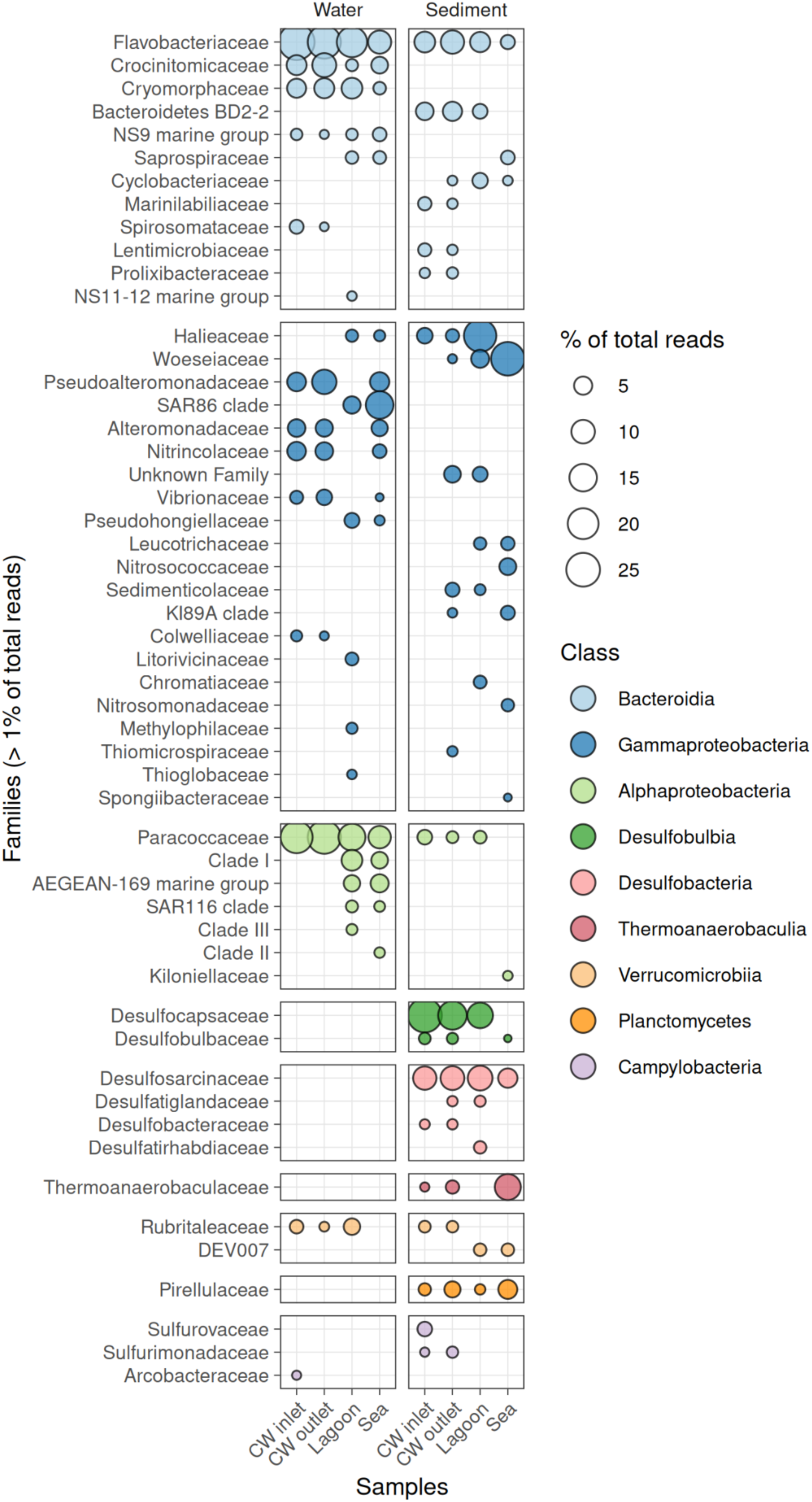
Relative abundance of dominant bacterial families aggregated by taxonomic class in water and sediment samples. Families with relative abundances greater than 1% of the total reads in at least one sample, and only the classes represented in at least three distinct samples, were included in the visualization. Class boxes were ordered by total class abundance, with the most abundant one at the top. Bubbles were sized proportionally to the relative abundance and colored according to the corresponding class.

The functional annotation of retrieved taxa through FAPROTAX showed the aerobic and anaerobic chemoheterotrophic processes were dominant, particularly in waters at the CW inlet and outlet. Regarding the nitrogen metabolism, nitrate reduction showed a relatively higher occurrence in water compared to the sediment samples. Conversely, the respiration of sulfate and sulfur compounds showed relatively higher values in sediments (Supplementary Figure 1). By interrogating the amplicon dataset using the FAPROTAX functions related to pathogenicity, a number of potential pathogenic taxa were identified. In both water and sediment samples, the CW inlet showed higher absolute values compared to the CW outlet. Human-associated pathogens and intracellular parasites were present mostly in the seawater sample (Figure 3a). The most represented genera, describing almost 60% of the abundances of the entire dataset, were unclassified at the genus level but mostly belonged to the order Rickettsiales. The seawater and the CW inlet contained the majority of the potential pathogens and parasites. The most represented classified genera were *Francisella*, *Campylobacter*, *Sphingomonas*, *Coxiella*, *Leifsonia*, and *Mycobacterium* (Figure 3b). The retrieved genera were present in the reference databases of medically relevant pathogenic bacteria, with MD3-55 as the only exception (Supplementary Table 3).

**Figure 3.**
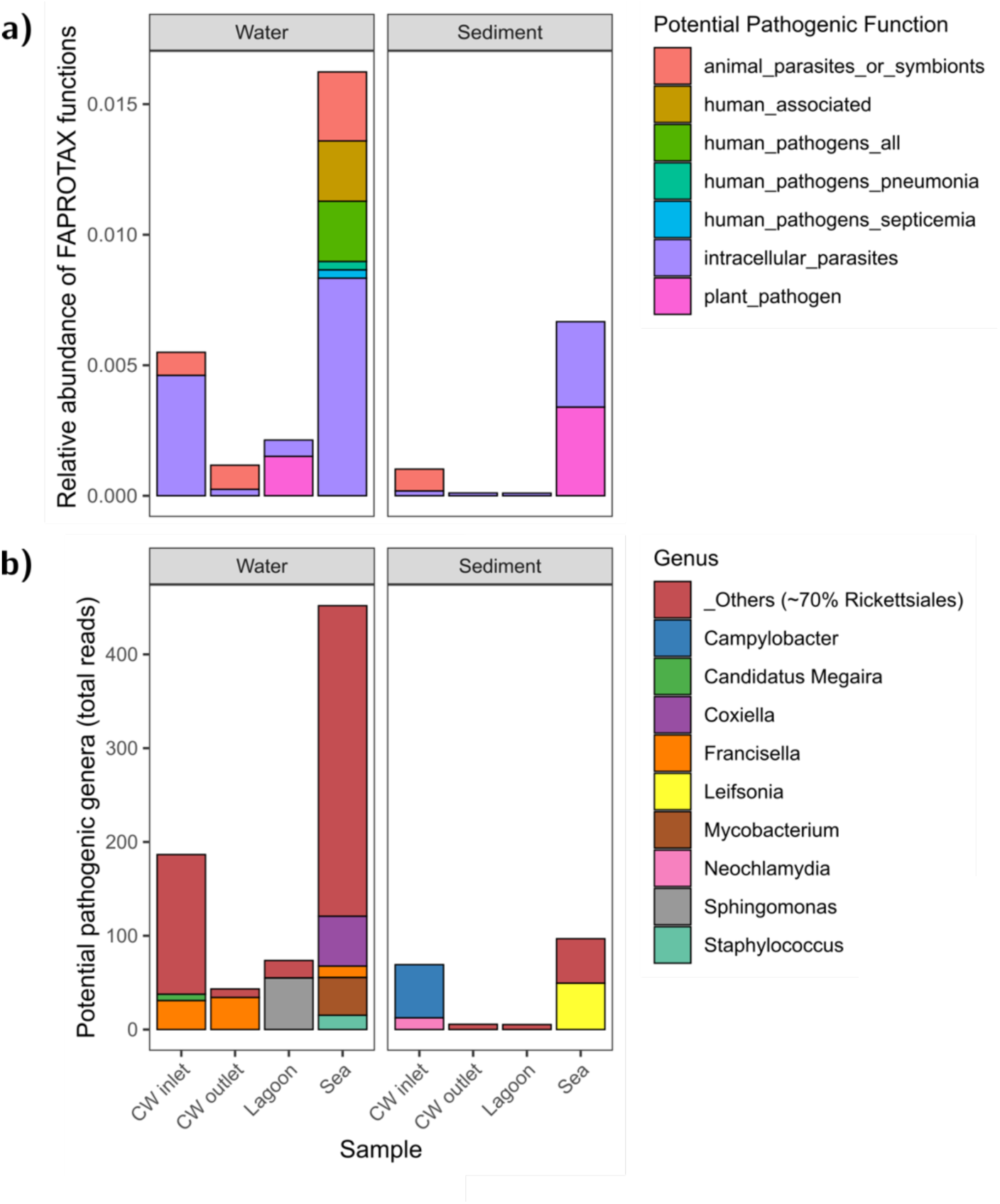
Pathogenicity functions associated with humans, animals, and plants across water and sediment samples (a). Total reads of top 10 potential pathogenic genera retrieved from the annotated functions (b).

### 3.3 Antimicrobial resistance genes and functional profiles from shotgun metagenomics

The highest occurrence of ARGs for each antimicrobial class category was found in seawater and sediment samples. Sediments showed a relatively higher occurrence of ARGs than water samples (Kruskal-Wallis test, p = 0.02). The richness of total ARGs was slightly higher in water than sediment, but the difference was not statistically significant (114.5 ± 10.0 and 93.8 ± 12.8 resistance genes per sample, respectively). The resistance to aminoglycosides was the most abundant, with an average of 11.4 ± 1.7 resistance genes per sample, followed by multidrug and fluoroquinolone/tetracycline resistance. The resistance genes to macrolides showed very low levels and were not retrieved in lagoon samples (Figure 4).

**Figure 4.**
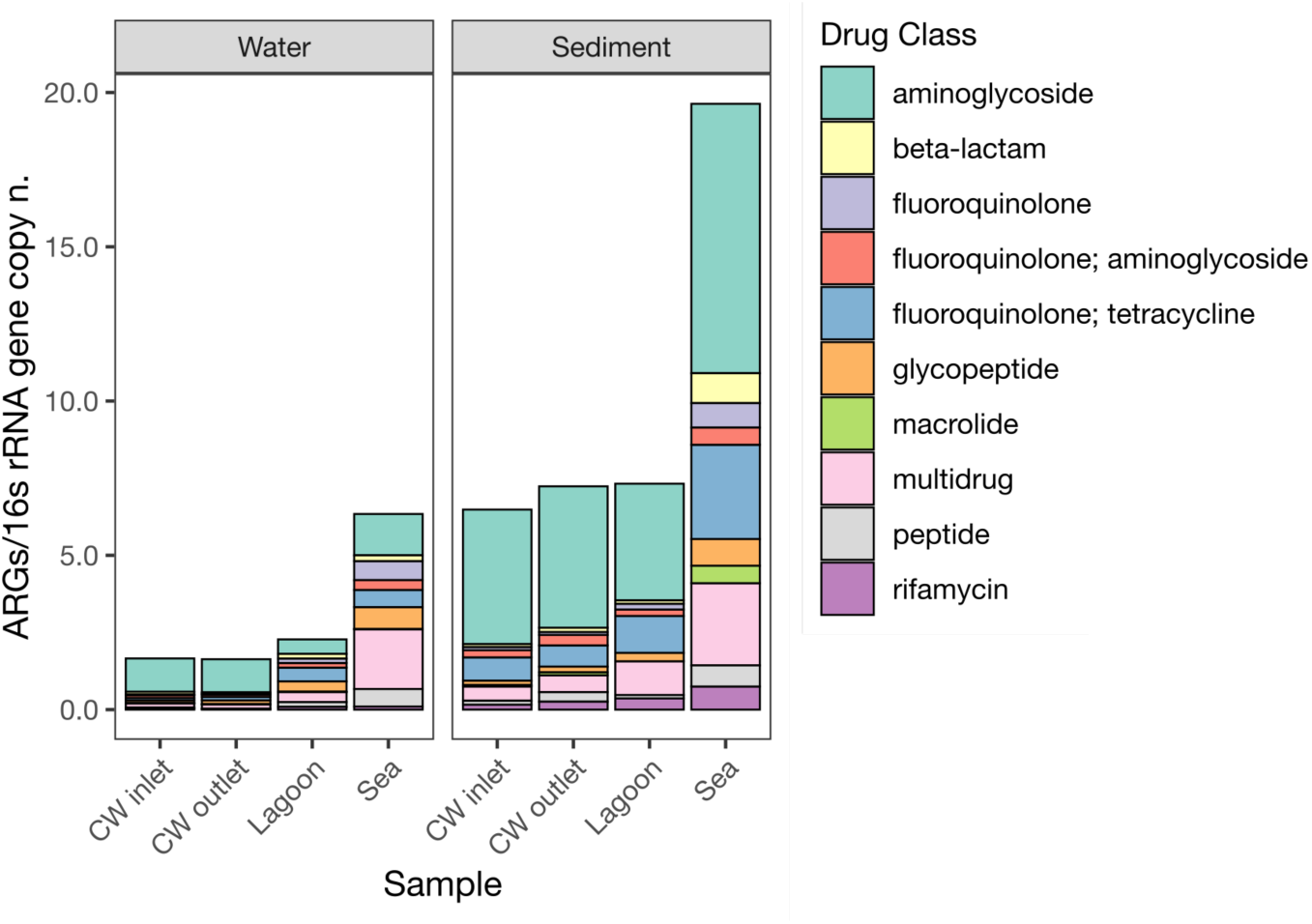
Occurrence of the top 10 antimicrobial resistance genes to drug classes. Gene copies were normalized by the 16S rRNA gene copy number. ARGs genes were identified from metagenomic data and processed using the RGI-bwt tool.

The functional annotation of metagenomic assemblies using DRAM supported the microbial preference for chemoheterotrophic processes (Supplementary Figure 3). Denitrification-related functions were the most represented in either water or sediment samples. Only three functions related to denitrification (i.e., nitrate reduction to nitrite and nitrite reduction to nitric oxide) and sulfur oxidation (i.e., thiosulfate oxidation to sulfate) were found in all samples. Overall, the CW inlet and outlet showed similar functional profiles and a consistent presence of the same functions, along with the environmental samples (Table 3; Supplementary Figure 4).

**Table 3.**
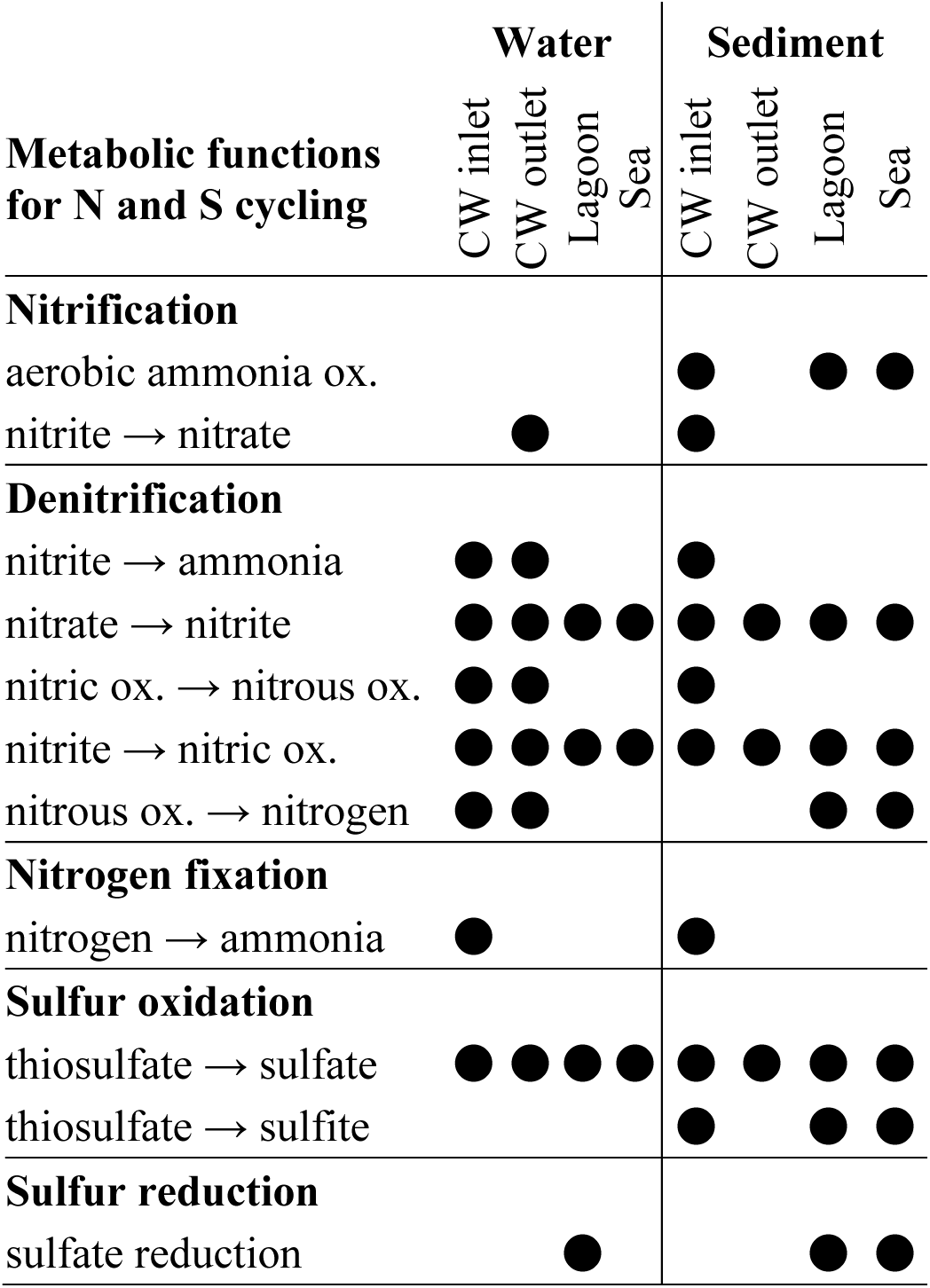
Presence of metabolic functions for nitrogen and sulfur cycles, retrieved with the functional annotation and distillation of DRAM on the corresponding shotgun assemblies.

### 3.4 Microbial metabolic potential

The average metabolic potential was similar between water and sediment (Figure 5).

**Figure 5.**
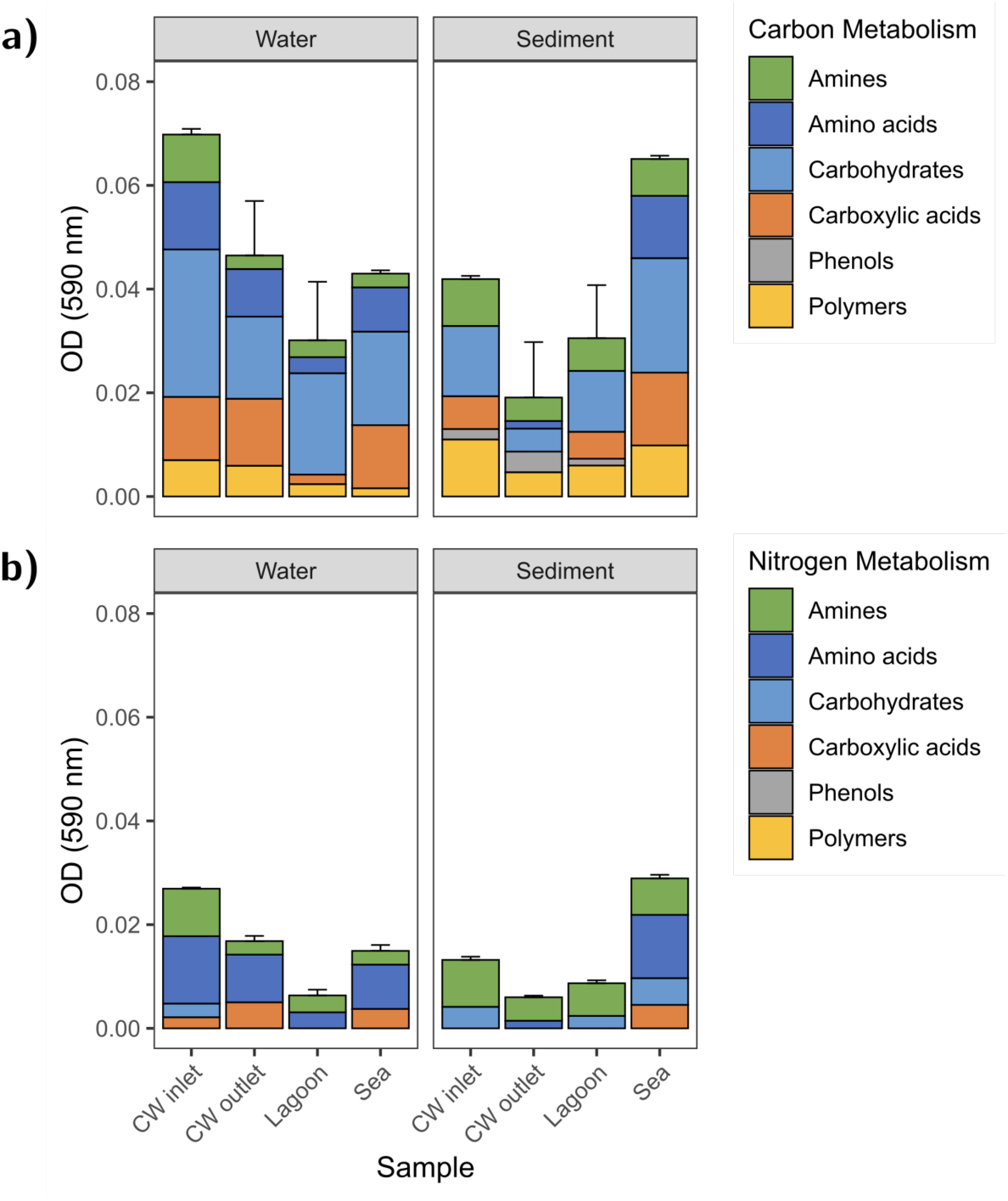
Average degradative activity assessed by the Biolog™ EcoPlates assay. Absorbance values (OD 590 nm) are proportional to the mineralization of 31 carbon sources (a) and 12 nitrogen sources (b), belonging to six classes of organic compounds (i.e., amines, amino acids, carbohydrates, carboxylic acids, phenols, and polymers).

The multivariate analysis did not show differences in C-metabolism, whereas N-containing substrates were differently metabolized (PERMANOVA test, p = 0.0053). More specifically, amino acids and carboxylic acids were preferentially degraded in water (Kruskal–Wallis test, p = 0.01 and p = 0.04, respectively), while amines and carbohydrates were preferentially degraded in sediments (Kruskal–Wallis test, p = 0.01 and p = 0.02, respectively). Water and sediments were significantly more active in C- and N-metabolic potential at the CW inlet than the outlet (Kruskal-Wallis test, p = 0.04). Moreover, N-enriched carbohydrates and amines were more degraded in sediments at the CW inlet than the outlet (Kruskal-Wallis test, p = 0.03 and p = 0.04, respectively). Notably, phenol degradation was not observed in water samples and very limited in sediment samples.

### 3.5 Multivariate statistics for data integration

The NMDS ordination biplot showed the difference in structural and functional properties of the aquatic microbial communities with an excellent fit (stress value = 0.053). The primary separation took place along the first NMDS axis, which significantly differentiated water from sediment samples (PERMANOVA, p = 0.03). The second axis represented the contrast between samples from CW and the natural environments, but the microbial community profiles were not statistically different (CW vs sea and lagoon, p > 0.05). The wider dispersion of water samples in the ordination space suggested a higher microbial heterogeneity than that found in sediment samples. Notably, the CW outlet was closer and more similar to the lagoon control samples than the CW inlet. Vectors representing dominant microbial taxa and metabolic traits varied markedly between sample types. Marine sediments were likely the most human-impacted samples, showing a higher concentration of potential pathogens and ARGs than any other sample. The vectors of family-level taxa were differently oriented among sample types (waters vs sediments), indicating stronger associations with the local environment (Figure 6).

**Figure 6.**
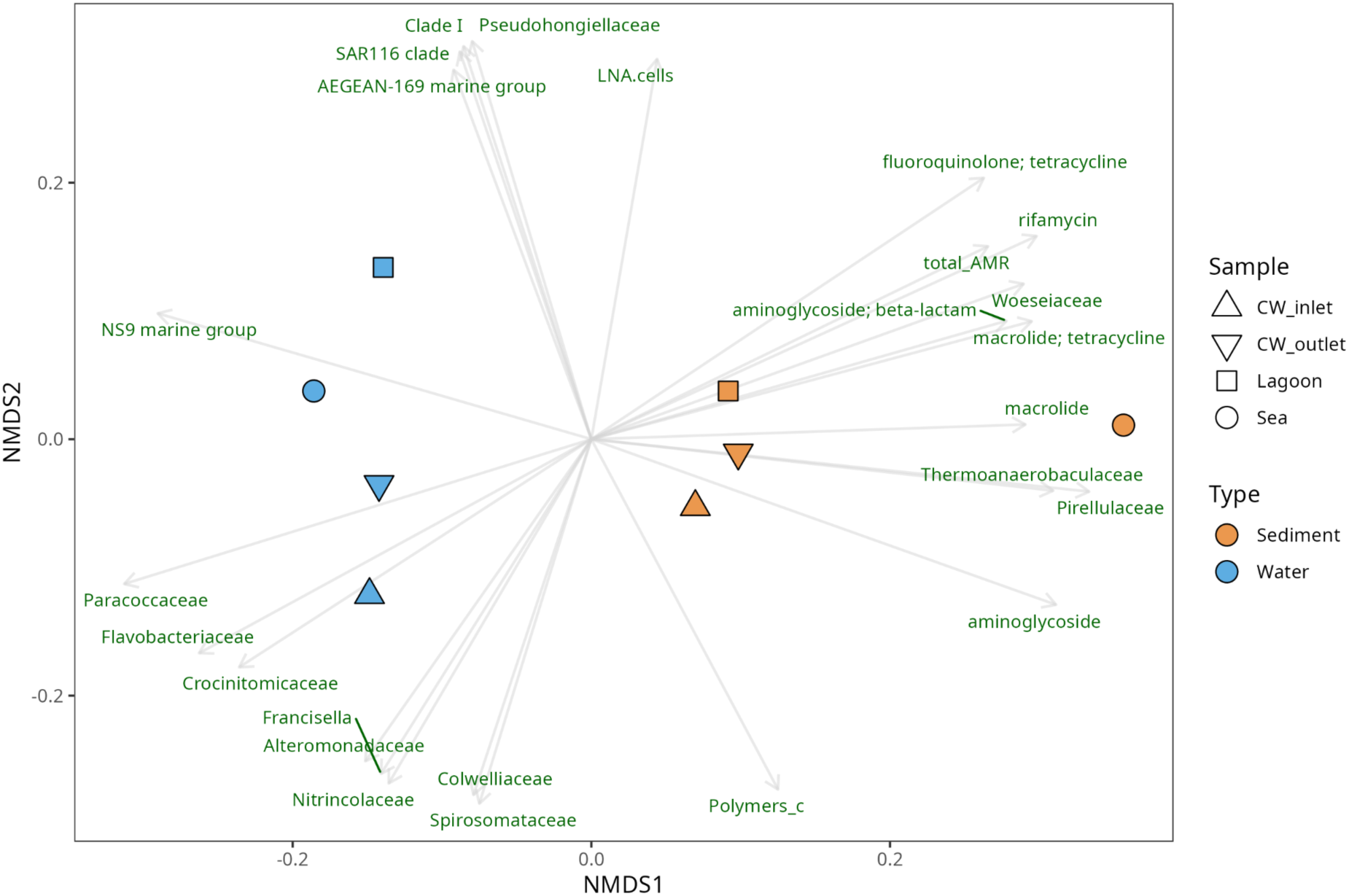
NMDS ordination integrating multiple data, such as flow cytometry counts, microbial community composition with phylogenetic Shannon index at the family level, occurrence of top ten potential pathogens and ARGs, and microbial functional profiles from Biolog assays (average degradative activity of carbon and nitrogen sources with functional Shannon index). Vectors represent variables significantly associated with the ordination (p < 0.05), scaled for best visualization.

## 4. Discussion

This study provided original field-based insights into significant microbial changes along the aquaculture–wetland–coastal continuum. While the CW effectively modulated microbial structure, limiting the spread of potential pathogens and ARGs, the microbial diversity and functional profiles remained relatively stable, pointing to a system that supports microbial resilience and gradual ecological restoration (Hollstein et al., 2023; Zhao et al., 2024). CWs are acknowledged to promote complex biogeochemical interactions and mitigate the ecological impact of aquaculture wastewater before discharge into natural environments (Xu et al., 2024). Unlike conventional treatment systems, CWs rely on the self-organizing properties of planktonic, benthic, and riparian communities, which together regulate nutrient dynamics, reduce pollutant loads, and modulate microbiological contamination risks (Kumar et al., 2023).

The target CW system presented a unique environment characterized by physical-chemical and microbial signatures reflecting the origin and treatment of aquaculture effluents with high organic and nutrient loads, a typical feature of aquaculture effluents (Boyd, 2003; Islam, 2005). While pH, salinity, and conductivity remained relatively stable along the CW flowpath, the differences with lagoon and marine controls confirmed that CW can create a transitional zone between the aquaculture-affected systems and the natural environments. The over-oxygenation observed at the CW outlet and lagoon sites suggested high photosynthetic activity, potentially stimulated by phytoplankton proliferation and oxygenation by emergent macrophytes (Vymazal et al., 2021).

The higher microbial load in sediments compared to the water column, especially in CW samples, aligns with the understanding that CW sediments can act as microbial hotspots, where organic matter accumulates and microbial activity intensifies (Mustafa et al., 2024; Yang et al., 2021). The dominance of HNA cells further supported the hypothesis that CW can promote metabolically active microbial communities in response to nutrient inputs (Amalfitano et al., 2018; Bouvier et al., 2007), suggesting a shift toward more diverse or functionally active assemblages compared to environmental controls.

The 16S rRNA gene amplicon sequencing revealed phylogenetic differences in microbial community composition primarily between water and sediment samples. This water–sediment dichotomy is a recurrent pattern in natural and engineered aquatic systems, generally attributed to different redox conditions, substrate availability, and dominant microbial processes (Fang et al., 2021; Wang et al., 2024). However, no significant difference emerged between CW and natural environmental samples in terms of alpha diversity or overall community composition, suggesting that CWs may facilitate the convergence of phylogenetic diversity under consistent environmental constraints (Grandmont-Lemire et al., 2025; Tian et al., 2024). The dominant taxa found in CW waters, including members of classes Bacteroidia (i.e., families Flavobacteriaceae, Crocinitomicaceae, Cryomorphaceae) and Alphaproteobacteria (i.e., family Paracoccaceae), are known to thrive in nutrient-rich aquatic environments and play key roles in the degradation of complex organic matter (Bowman, 2020; Moschos et al., 2022; Sun et al., 2012). In contrast, CW sediments were dominated by Thermodesulfobacteriota (i.e., families Desulfocapsaceae and Desulfosarcinaceae), known for their roles in anaerobic sulfate reduction and organic matter mineralization under anoxic conditions (Anantharaman et al., 2018). These findings indicated the establishment of functionally specialized communities, wherein microorganisms can be adapted for shorter-term organic matter turnover in water and for longer-term nutrient cycling in sediments (Brailsford et al., 2019; Wang et al., 2022).

Our results also confirmed the role of CWs in reducing the overall pathogen load by acting as nature-based barriers, primarily due to natural processes (Borsetto et al., 2025; Sleytr et al., 2007; Wu et al., 2016). The reduced abundance of potentially pathogenic taxa from the CW inlet to the outlet, including genera such as *Francisella* and *Campylobacter*, suggested a partial attenuation of microbiological hazards. The validation of the retrieved potential pathogens across different databases indicated that the majority of identified genera are well-documented and may contain species with pathogenic traits, although some genera are still poorly characterized (i.e., MD3-55). The persistence of some pathogenic genera in seawater samples pointed to a cumulative impact from multiple pollution sources. The distribution patterns of ARGs also support these conclusions. The highest ARG frequencies were observed in marine sediments, since various anthropogenic sources may converge in coastal depositional zones (Wanyan et al., 2023). Previous studies noted that sediments may serve as reservoirs for allochthonous and antimicrobial-resistant bacteria because they provide a more protective environment compared to water regarding environmental stressors (Byappanahalli et al., 2012; Di Cesare et al., 2014). The predominance of resistance genes to aminoglycosides, fluoroquinolones, and tetracyclines reflected the common use of these antimicrobials in anthropogenic activities and human medicine (Yang et al., 2013). Although the ARG abundance showed particularly lower values compared to other wastewater-fed CWs (Ma et al., 2022), the persistence patterns of key resistance genes indicated that CWs may serve as partial buffers against the release of ARGs into surrounding ecosystems (Borsetto et al., 2025; Sabri et al., 2021; Zhang et al., 2025). While the CW system did not eliminate antimicrobial resistance determinants, it did not appear to amplify them either, further highlighting the efficiency of CWs in controlling the microbial contamination spread (Ma et al., 2022).

It is also important to consider the role of local wildlife in introducing microbiological contaminants of health concern. Seagulls and other aquatic birds are known to shed enteric bacteria, protozoan parasites, and associated ARGs into wetlands and adjacent marine environments, acting as both local and migratory vectors (Jarma et al., 2024; Skarżyńska et al., 2021). Similarly, terrestrial mammals (i.e., rodents, cats, foxes, dogs, and wild boars were sighted during the sampling period) may contribute to fecal contamination in ecotone areas where CWs interface with natural wetlands. Such wildlife-mediated pathways may help explain the persistence of microbiological contaminants in CWs (Collins, 2004; Cookson et al., 2024; Li et al., 2021).

The functional annotations derived from FAPROTAX and DRAM analyses converged in identifying key microbial pathways associated with the transformation of carbon, nitrogen, and sulfur compounds. The dominant chemoheterotrophic metabolism in both water and sediment samples is likely to reflect the high input of organic substrates and nutrients typically found in fish farming effluents (Moschos et al., 2022; Truu et al., 2009). This metabolic profile remained consistently high at both the CW inlet and outlet, suggesting that microbial communities continue to be active in processing organic matter along the CW pathway. Nitrogen metabolism was mainly characterized by denitrification-related processes, particularly nitrate and nitrite reduction. Our findings confirmed the known role of CWs in promoting nitrogen removal through denitrification, especially under low oxygen and high carbon levels (Ilyas and Masih, 2017; Kamilya et al., 2022). In sediments, sulfur-respiring functions were also likely to occur, reflecting anaerobic conditions where sulfate reducers thrive (Liu et al., 2021; Wu et al., 2013). Importantly, the functions detected across all sample types (nitrate reduction to nitrite, nitrite reduction to nitric oxide, and thiosulfate oxidation to sulfate) underscored a baseline of functional redundancy, confirming that microbial communities in CWs can maintain essential metabolic capacities for nutrient cycling (Allison and Martiny, 2008; Zhao et al., 2024). The BIOLOG EcoPlate assays confirmed that microbial communities retained the capacity to degrade a broad spectrum of carbon and nitrogen sources. Amino acids and carbohydrates were the most efficiently degraded substrates as carbon and nitrogen sources, while phenols and polymers were more recalcitrant (Rana et al., 2021; Salomo et al., 2009). The similar average degradative activity between water and sediment samples suggested a functionally resilient community, able to maintain metabolic activity across environmental gradients (Griffiths and Philippot, 2013; Hollstein et al., 2023; Tian et al., 2024). The CW inlet showed higher metabolic activity than the outlet, likely due to the inputs of labile organic matter from aquaculture effluents. However, despite the observed shifts, the metabolic diversity remained relatively high, and the functional profiles were not drastically reshaped. This evidence is likely to support the notion that CWs can select for functionally competent microbial assemblages capable of sustaining essential ecosystem services even under anthropogenic pressure (Rani et al., 2024; Zhao et al., 2024).

## 5. Conclusion

Our findings demonstrate that CWs can significantly influence the structure and function of aquatic microbial communities associated with aquaculture wastewater. CWs promoted microbial attenuation, reduced the occurrence of potentially pathogenic taxa, and modulated the distribution of ARGs while promoting nutrient removal (i.e., denitrification) without compromising major biogeochemical cycles and essential ecosystem functions. Nonetheless, CWs did not eliminate potential microbiological hazards, with sediments representing a potential reservoir of microbial and genetic signatures. The combined influence of aquaculture discharges and natural wildlife-mediated inputs must also be considered when interpreting pathogen and antimicrobial resistance dynamics in CW systems. By employing a multi-omics approach and ecological baselines, our findings emphasize the importance of integrated control strategies, such as extending hydraulic retention times, promoting diverse vegetation, or incorporating complementary treatment stages for aquaculture effluent management. From a broader perspective, this study contributes to the understanding of nature-based solutions by providing field insights that demonstrate how CWs support the transition from high-impact effluents to ecologically sustainable reclamation options.

## Acknowledgments

The authors would like to thank the EU and the Ministry of University and Research (MUR, Italy) for funding in the frame of the collaborative international consortium ARENA financed under the ERA-NET AquaticPollutants Joint Transnational Call (GA n. 869178). This work was also funded by the project “National Biodiversity Future Center - NBFC” under the National Recovery and Resilience Plan (NRRP). This study represents partial fulfillment of the requirements for the PhD thesis of Naomi Massaccesi at the FishMed-PhD course (https://www.FishMed-PhD.org). We thank the fish farm personnel for their kind support and for granting access to the aquaculture facilities.

## 5. Code Availability

All the codes used for the 16S rRNA gene analysis can be retrieved in the following GitHub Repository (https://github.com/MicroEcoLab/CW-Orbetello).

## Supplementary material

**Supplementary Table 1.** The sequencing data can be retrieved for each sample by using the BioProject PRJNA1200612 and the corresponding sample ID from the NCBI database.

| Sample | Amplicon sequences<br>(16S rRNA genes) |  | Shotgun whole genome sequences |  |
| --- | --- | --- | --- | --- |
|  | Water | Sediment | Water | Sediment |
| CW inlet | SAMN45887343 | SAMN45887350 | SAMN46716068 | SAMN46716082 |
| CW outlet | SAMN45887344 | SAMN45887351 | SAMN46716076 | SAMN46716083 |
| Lagoon | SAMN45887345 | SAMN45887354 | SAMN46716070 | SAMN46716086 |
| Sea | SAMN45887346 | SAMN45887353 | SAMN46716078 | SAMN46716085 |

**Supplementary Table 2.**
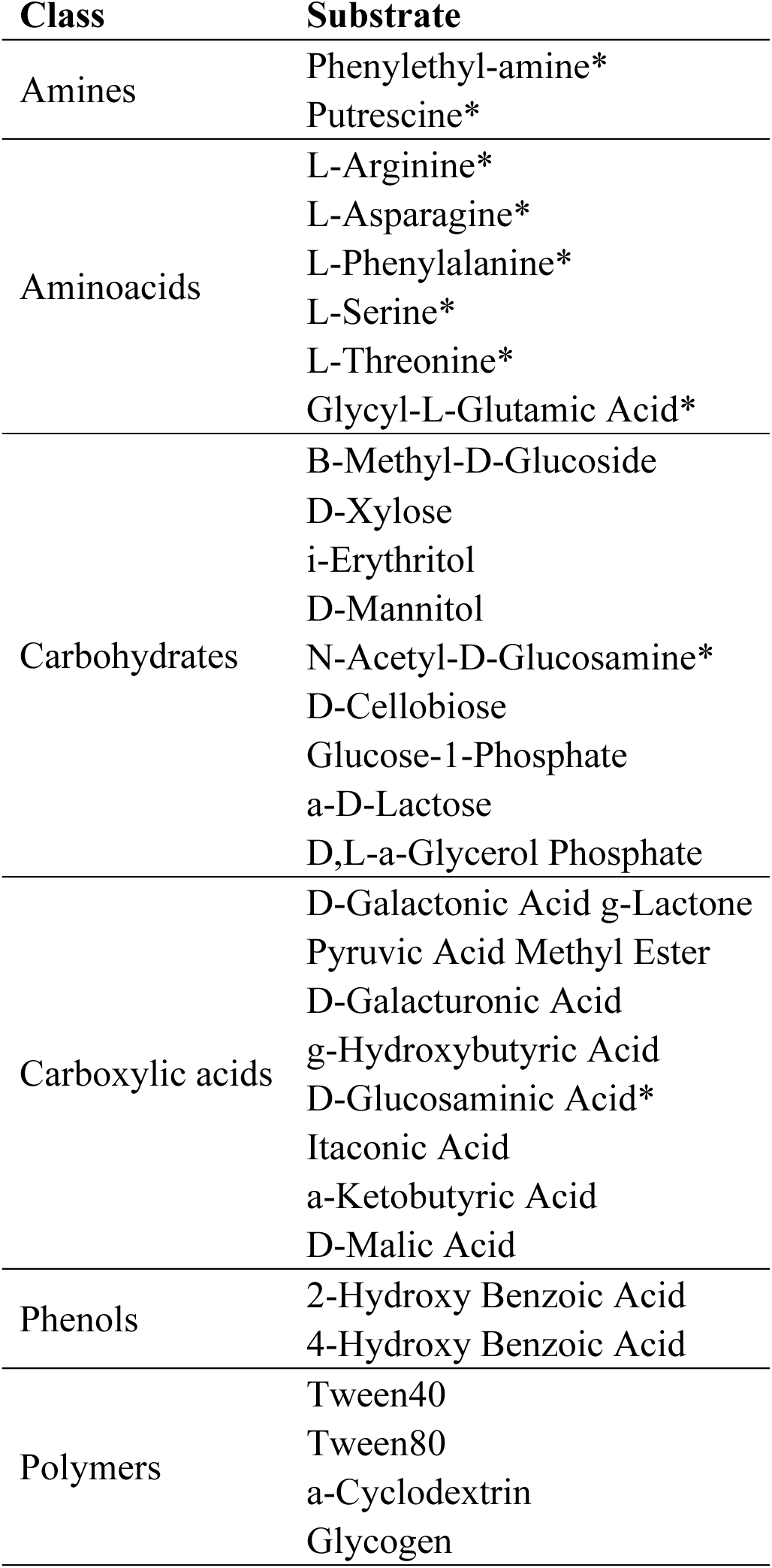
Substrates of the BIOLOG EcoPlates grouped by macroclass of organic compounds. Asterisks indicate N-containing compounds.

**Supplementary Table 3.**
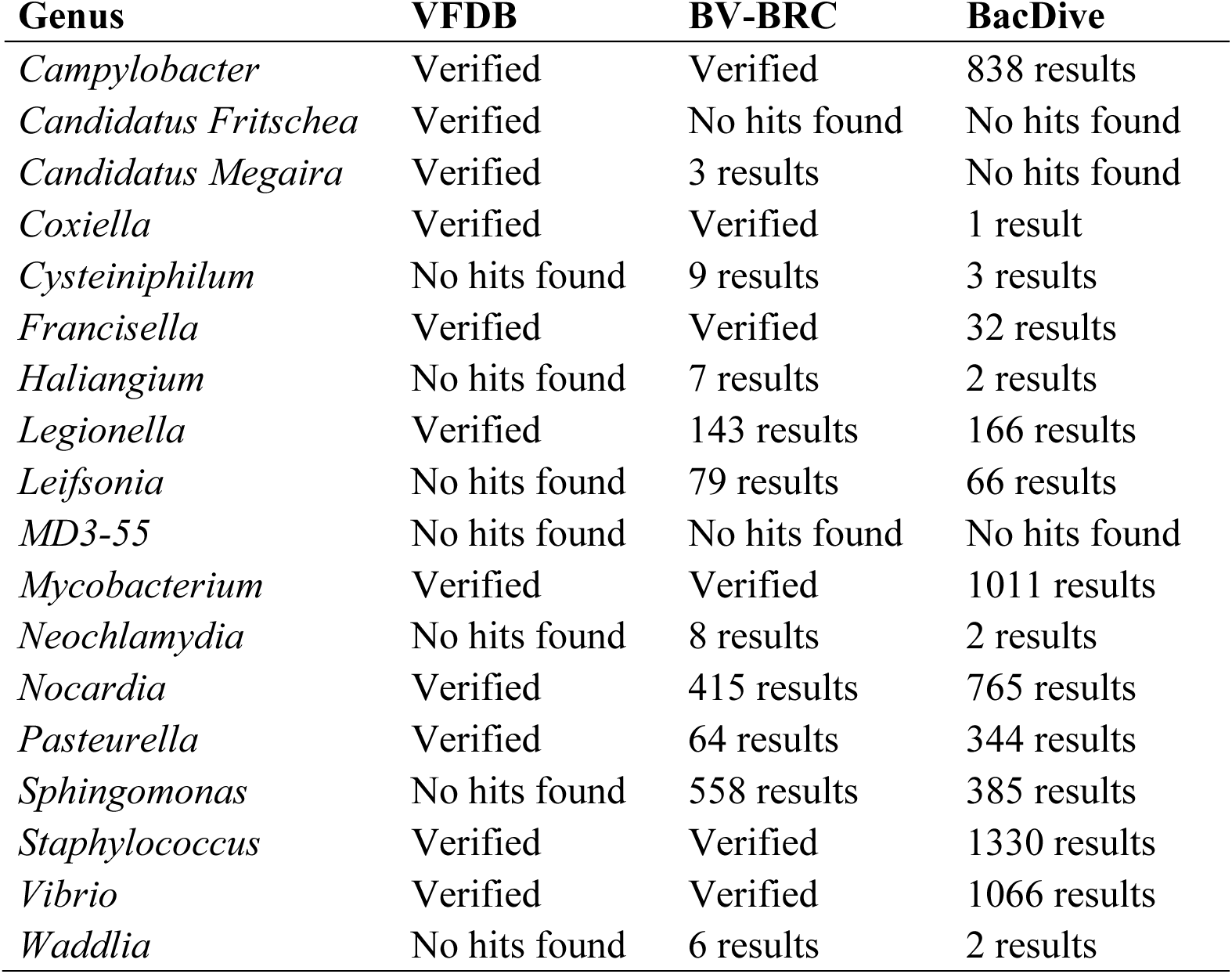
Validation of genera associated with potential pathogenicity (as identified by FAPROTAX functions) across three reference databases: the Virulence Factors Database (VFDB), the Bacterial and Viral Bioinformatics Resource Center (BV-BRC, search by taxa), and BacDive (search by genus).

**Supplementary Figure 1.**
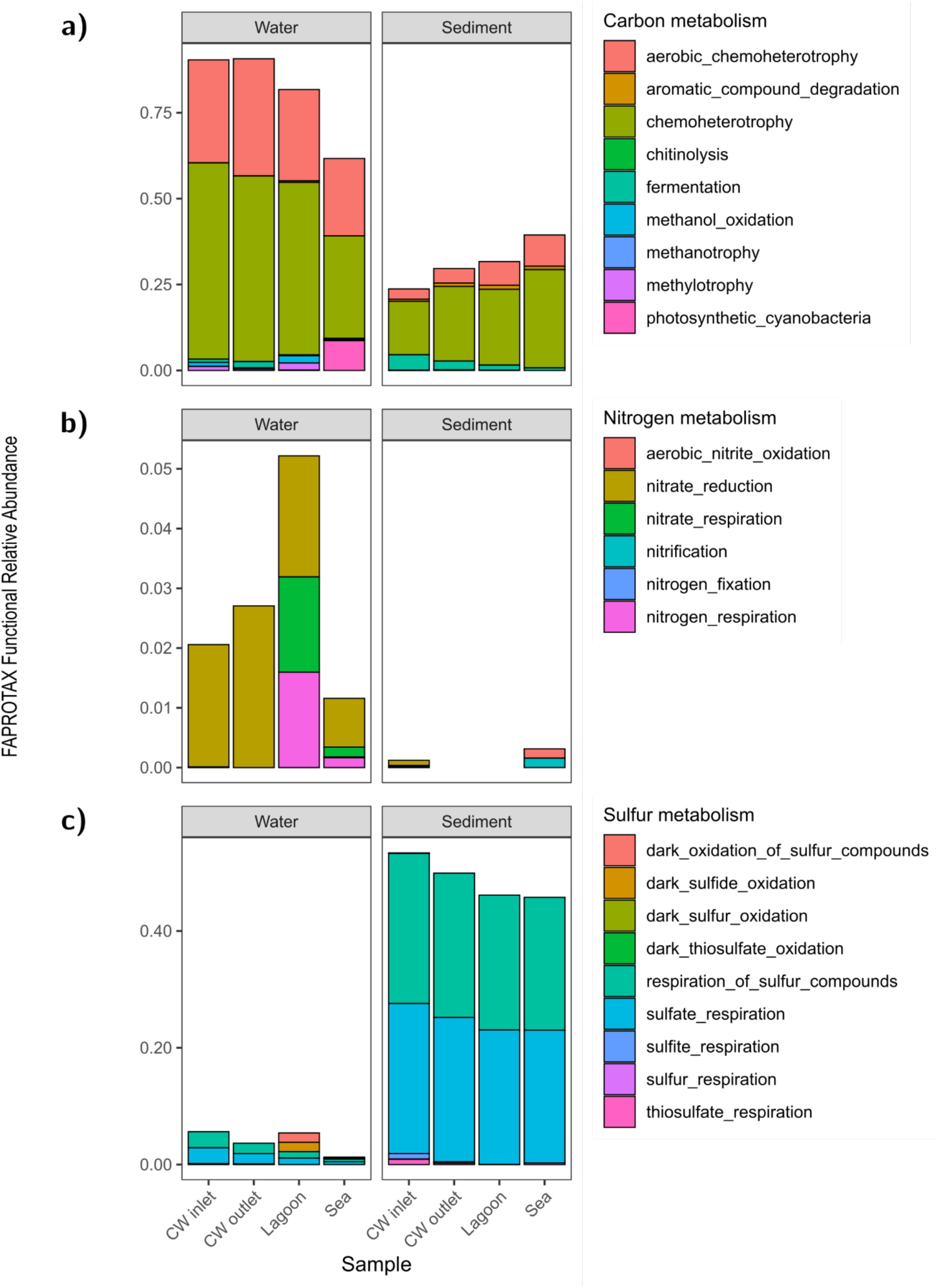
Putative metabolic functions of microbial taxa annotated by FAPROTAX. Barplots show the relative abundance of functions related to carbon (a), nitrogen (b), and sulfur (c).

**Supplementary Figure 2.**
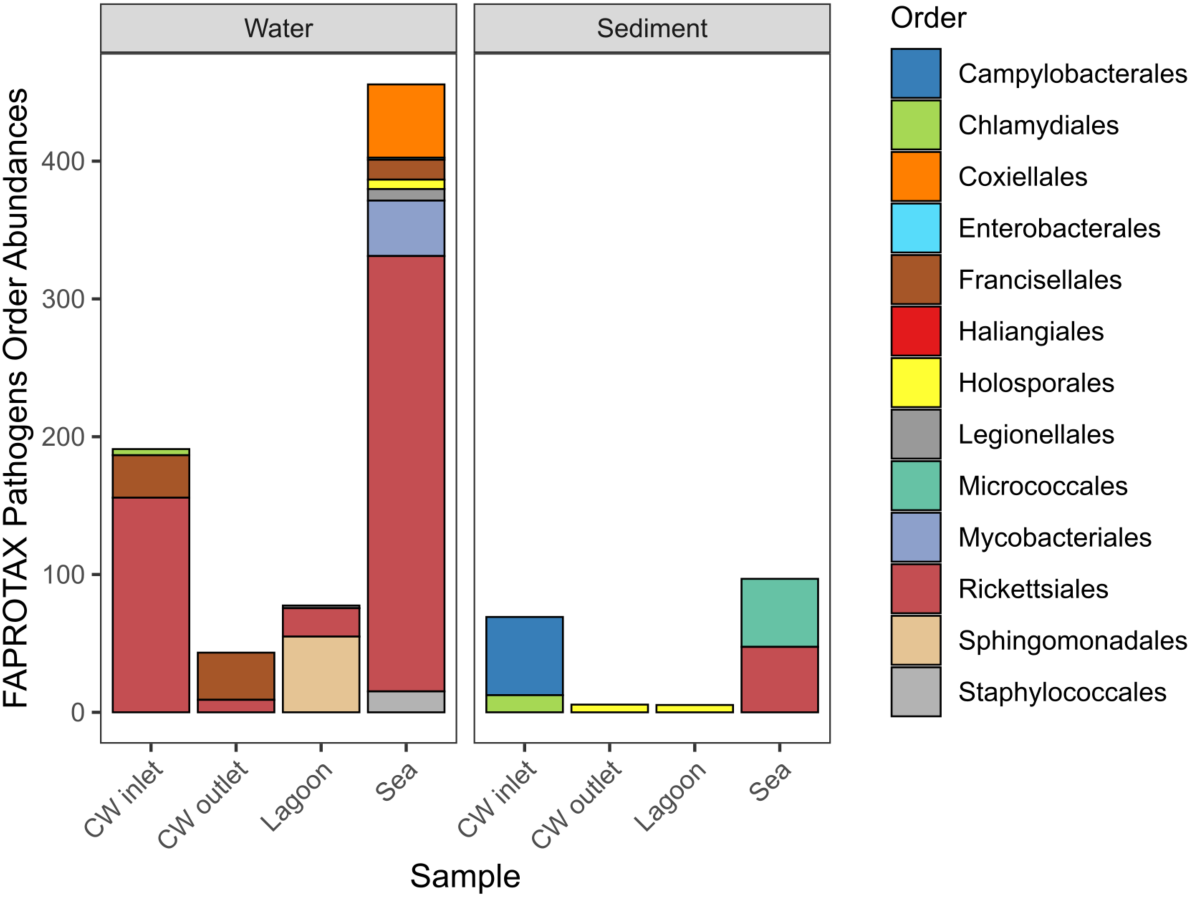
Functional abundances of bacterial taxa at the order level associated with potential pathogenic functions predicted by FAPROTAX across different sampling sites.

**Supplementary Figure 3.**
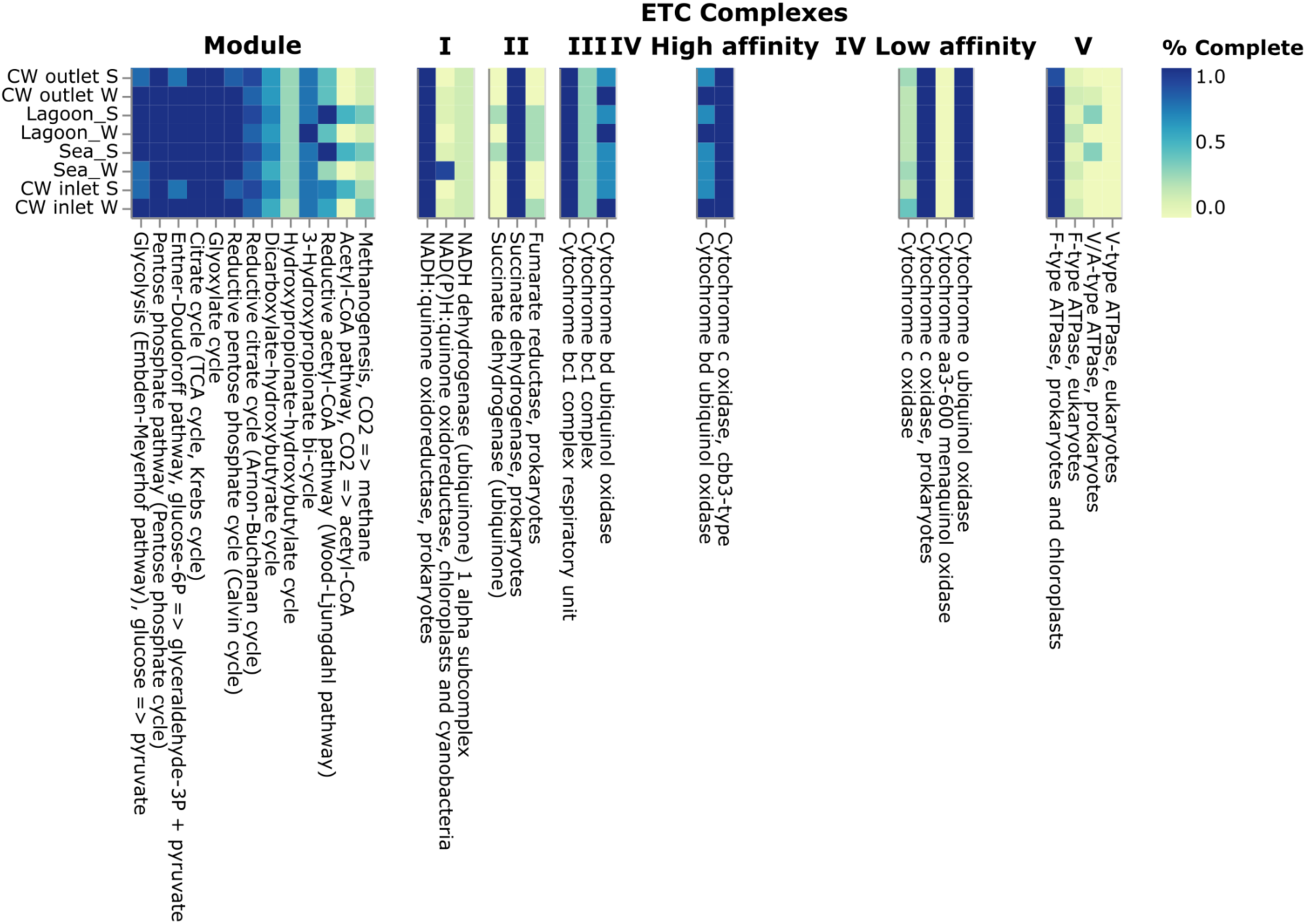
DRAM heatmap output showing the presence and completeness of electron transport chain complexes identified in corresponding sample assemblies obtained from metagenomics data. Color intensity indicates completeness, with darker shades representing higher completeness.

**Supplementary figure 4.**
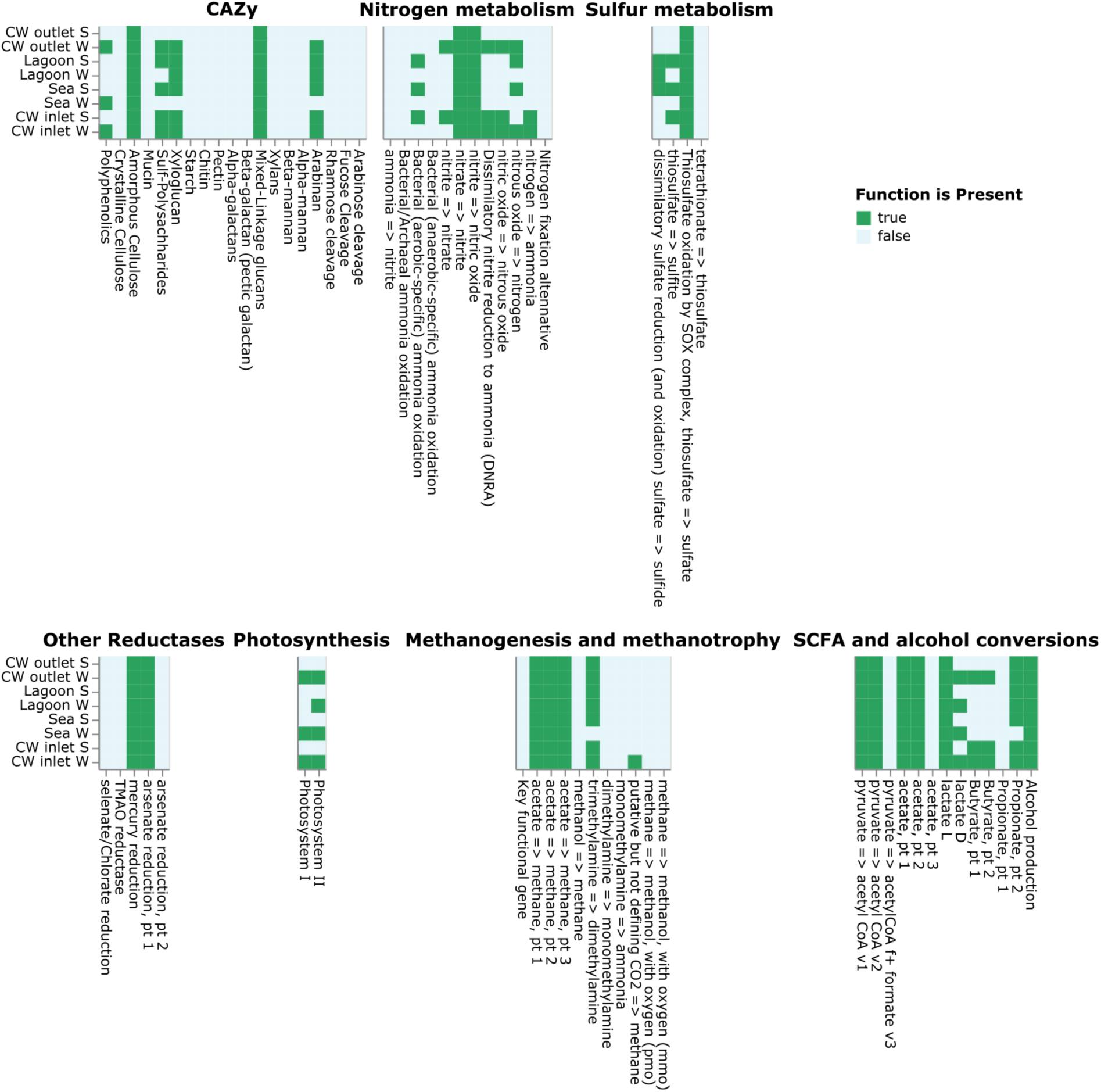
DRAM heatmap output illustrating the presence or absence of major metabolic functions across obtained assemblies as determined by DRAM. Metabolisms include CAZy, nitrogen, sulfur, reductases, photosynthesis, methanogenesis and methanotrophy, and SCFA and alcohol conversions.

## References

Ahmad, A.L., Chin, J.Y., Mohd Harun, M.H.Z., Low, S.C., 2022. Environmental impacts and imperative technologies towards sustainable treatment of aquaculture wastewater: A review. Journal of Water Process Engineering 46, 102553. 10.1016/j.jwpe.2021.102553

Alcock, B.P., Huynh, W., Chalil, R., Smith, K.W., Raphenya, A.R., Wlodarski, M.A., Edalatmand, A., Petkau, A., Syed, S.A., Tsang, K.K., Baker, S.J.C., Dave, M., McCarthy, M.C., Mukiri, K.M., Nasir, J.A., Golbon, B., Imtiaz, H., Jiang, X., Kaur, K., Kwong, M., Liang, Z.C., Niu, K.C., Shan, P., Yang, J.Y.J., Gray, K.L., Hoad, G.R., Jia, B., Bhando, T., Carfrae, L.A., Farha, M.A., French, S., Gordzevich, R., Rachwalski, K., Tu, M.M., Bordeleau, E., Dooley, D., Griffiths, E., Zubyk, H.L., Brown, E.D., Maguire, F., Beiko, R.G., Hsiao, W.W.L., Brinkman, F.S.L., Van Domselaar, G., McArthur, A.G., 2023. CARD 2023: expanded curation, support for machine learning, and resistome prediction at the Comprehensive Antibiotic Resistance Database. Nucleic Acids Research 51, D690–D699. 10.1093/nar/gkac920

Allison, S.D., Martiny, J.B.H., 2008. Resistance, resilience, and redundancy in microbial communities. Proceedings of the National Academy of Sciences 105, 11512–11519. 10.1073/pnas.0801925105

Amalfitano, S., Fazi, S., 2008. Recovery and quantification of bacterial cells associated with streambed sediments. Journal of Microbiological Methods 75, 237–243. 10.1016/j.mimet.2008.06.004

Amalfitano, S., Fazi, S., Ejarque, E., Freixa, A., Romaní, A.M., Butturini, A., 2018. Deconvolution model to resolve cytometric microbial community patterns in flowing waters. Cytometry Part A 93, 194–200. 10.1002/cyto.a.23304

Anantharaman, K., Hausmann, B., Jungbluth, S.P., Kantor, R.S., Lavy, A., Warren, L.A., Rappé, M.S., Pester, M., Loy, A., Thomas, B.C., Banfield, J.F., 2018. Expanded diversity of microbial groups that shape the dissimilatory sulfur cycle. ISME J 12, 1715–1728. 10.1038/s41396-018-0078-0

Andrews, S., 2010. FastQC: A quality control tool for high throughput sequence data.

Arkin, A.P., Cottingham, R.W., Henry, C.S., Harris, N.L., Stevens, R.L., Maslov, S., Dehal, P., Ware, D., Perez, F., Canon, S., Sneddon, M.W., Henderson, M.L., Riehl, W.J., Murphy-Olson, D., Chan, S.Y., Kamimura, R.T., Kumari, S., Drake, M.M., Brettin, T.S., Glass, E.M., Chivian, D., Gunter, D., Weston, D.J., Allen, B.H., Baumohl, J., Best, A.A., Bowen, B., Brenner, S.E., Bun, C.C., Chandonia, J.-M., Chia, J.-M., Colasanti, R., Conrad, N., Davis, J.J., Davison, B.H., DeJongh, M., Devoid, S., Dietrich, E., Dubchak, I., Edirisinghe, J.N., Fang, G., Faria, J.P., Frybarger, P.M., Gerlach, W., Gerstein, M., Greiner, A., Gurtowski, J., Haun, H.L., He, F., Jain, R., Joachimiak, M.P., Keegan, K.P., Kondo, S., Kumar, V., Land, M.L., Meyer, F., Mills, M., Novichkov, P.S., Oh, T., Olsen, G.J., Olson, R., Parrello, B., Pasternak, S., Pearson, E., Poon, S.S., Price, G.A., Ramakrishnan, S., Ranjan, P., Ronald, P.C., Schatz, M.C., Seaver, S.M.D., Shukla, M., Sutormin, R.A., Syed, M.H., Thomason, J., Tintle, N.L., Wang, D., Xia, F., Yoo, H., Yoo, S., Yu, D., 2018. KBase: The United States Department of Energy Systems Biology Knowledgebase. Nat Biotechnol 36, 566–569. 10.1038/nbt.4163

Barros, D.J., Carvalho, G.A., de Chaves, M.G., Vanzela, L.S., Kozusny-Andreani, D.I., Guarda, E.A., Neu, V., de Morais, P.B., Tsai, S.M., Navarrete, A.A., 2023. Microbial metabolic activity in Amazon floodplain forest and agricultural soils. Front. Microbiol. 14. 10.3389/fmicb.2023.1144062

Basili, M., Perini, L., Zaggia, L., Luna, G.M., Quero, G.M., 2023. Integrating culture-based and molecular methods provides an improved assessment of microbial quality in a coastal lagoon. Environmental Pollution 334, 122140. 10.1016/j.envpol.2023.122140

Bolger, A.M., Lohse, M., Usadel, B., 2014. Trimmomatic: a flexible trimmer for Illumina sequence data. Bioinformatics 30, 2114–2120. 10.1093/bioinformatics/btu170

Borsetto, C., Dykes, C., Kockiri, B., Song, L., Wellington, E.M., Abolfathi, S., 2025. Constructed wetlands as nature-based barriers: Mitigating antimicrobial resistance and pathogen dispersal in riverine systems. Journal of Hazardous Materials 495, 138855. 10.1016/j.jhazmat.2025.138855

Bouvier, T., Del Giorgio, P.A., Gasol, J.M., 2007. A comparative study of the cytometric characteristics of High and Low nucleic-acid bacterioplankton cells from different aquatic ecosystems. Environmental Microbiology 9, 2050–2066. 10.1111/j.1462-2920.2007.01321.x

Bowman, J.P., 2020. Out From the Shadows – Resolution of the Taxonomy of the Family Cryomorphaceae. Front. Microbiol. 11. 10.3389/fmicb.2020.00795

Boyd, C.E., 2003. Guidelines for aquaculture effluent management at the farm-level. Aquaculture, Management of Aquaculture Effluents 226, 101–112. 10.1016/S0044-8486(03)00471-X

Brailsford, F.L., Glanville, H.C., Golyshin, P.N., Johnes, P.J., Yates, C.A., Jones, D.L., 2019. Microbial uptake kinetics of dissolved organic carbon (DOC) compound groups from river water and sediments. Sci Rep 9, 11229. 10.1038/s41598-019-47749-6

Byappanahalli, M.N., Nevers, M.B., Korajkic, A., Staley, Z.R., Harwood, V.J., 2012. Enterococci in the Environment. Microbiology and Molecular Biology Reviews 76, 685– 706. 10.1128/mmbr.00023-12

Callahan, B.J., McMurdie, P.J., Rosen, M.J., Han, A.W., Johnson, A.J.A., Holmes, S.P., 2016. DADA2: High-resolution sample inference from Illumina amplicon data. Nat Methods 13, 581–583. 10.1038/nmeth.3869

Collins, R., 2004. Fecal Contamination of Pastoral Wetlands. Journal of Environmental Quality 33, 1912–1918. 10.2134/jeq2004.1912

Cookson, A.L., Marshall, J.C., Biggs, P.J., Rogers, L.E., Collis, R.M., Devane, M., Stott, R., Brightwell, G., 2024. Impact of land-use and fecal contamination on Escherichia populations in environmental samples. Sci Rep 14, 32099. 10.1038/s41598-024-83594-y

Di Cesare, A., Pasquaroli, S., Vignaroli, C., Paroncini, P., Luna, G.M., Manso, E., Biavasco, F., 2014. The marine environment as a reservoir of enterococci carrying resistance and virulence genes strongly associated with clinical strains. Environmental Microbiology Reports 6, 184–190. 10.1111/1758-2229.12125

Fang, G., Yu, H., Sheng, H., Tang, Y., Liang, Z., 2021. Comparative analysis of microbial communities between water and sediment in Laoshan Bay marine ranching with varied aquaculture activities. Marine Pollution Bulletin 173, 112990. 10.1016/j.marpolbul.2021.112990

Gasol, J.M., Morán, X.A.G., 2016. Flow Cytometric Determination of Microbial Abundances and Its Use to Obtain Indices of Community Structure and Relative Activity, in: McGenity, T.J., Timmis, K.N., Nogales, B. (Eds.), Hydrocarbon and Lipid Microbiology Protocols: Single-Cell and Single-Molecule Methods. Springer, Berlin, Heidelberg, pp. 159–187. 10.1007/8623_2015_139

Grandmont-Lemire, S., Gearheart, B., Cuellar-Gempeler, C., 2025. Environmental effects on constructed wetland microbial diversity and function in the context of wastewater management. 10.1101/2025.01.14.633069

Griffiths, B.S., Philippot, L., 2013. Insights into the resistance and resilience of the soil microbial community. FEMS Microbiol Rev 37, 112–129. 10.1111/j.1574-6976.2012.00343.x

Hassan, I., Chowdhury, S.R., Prihartato, P.K., Razzak, S.A., 2021. Wastewater Treatment Using Constructed Wetland: Current Trends and Future Potential. Processes 9, 1917. 10.3390/pr9111917

Hollstein, M., Comerford, M., Uhl, M., Abel, M., Egan, S.P., Stadler, L.B., 2023. Impact of a natural disturbance on the performance and microbial communities in a full-scale constructed wetland for industrial wastewater treatment. Front. Environ. Sci. 11. 10.3389/fenvs.2023.1187143

Ilyas, H., Masih, I., 2017. The performance of the intensified constructed wetlands for organic matter and nitrogen removal: A review. Journal of Environmental Management 198, 372–383. 10.1016/j.jenvman.2017.04.098

Islam, Md.S., 2005. Nitrogen and phosphorus budget in coastal and marine cage aquaculture and impacts of effluent loading on ecosystem: review and analysis towards model development. Marine Pollution Bulletin 50, 48–61. 10.1016/j.marpolbul.2004.08.008

Jarma, D., Sacristán-Soriano, O., Borrego, C.M., Hortas, F., Peralta-Sánchez, J.M., Balcázar, J.L., Green, A.J., Alonso, E., Sánchez-Melsió, A., Sánchez, M.I., 2024. Variability of faecal microbiota and antibiotic resistance genes in flocks of migratory gulls and comparison with the surrounding environment. Environmental Pollution 359, 124563. 10.1016/j.envpol.2024.124563

Kamilya, T., Majumder, A., Yadav, M.K., Ayoob, S., Tripathy, S., Gupta, A.K., 2022. Nutrient pollution and its remediation using constructed wetlands: Insights into removal and recovery mechanisms, modifications and sustainable aspects. Journal of Environmental Chemical Engineering 10, 107444. 10.1016/j.jece.2022.107444

Kumar, V., Bera, T., Roy, S., Vuong, P., Jana, C., Sarkar, D.J., Devi, M.S., Jana, A.K., Rout, A.K., Kaur, P., Das, B.K., Behera, B.K., 2023. Investigating bio-remediation capabilities of a constructed wetland through spatial successional study of the sediment microbiome. npj Clean Water 6, 8. 10.1038/s41545-023-00225-1

Kushwaha, A., Goswami, L., Kim, B.S., Lee, S.S., Pandey, S.K., Kim, K.-H., 2024. Constructed wetlands for the removal of organic micropollutants from wastewater: Current status, progress, and challenges. Chemosphere 360, 142364. 10.1016/j.chemosphere.2024.142364

Li, Q., Long, Z., Wang, H., Zhang, G., 2021. Functions of constructed wetland animals in water environment protection – A critical review. Science of The Total Environment 760, 144038. 10.1016/j.scitotenv.2020.144038

Liu, W., Rahaman, Md.H., Mąkinia, J., Zhai, J., 2021. Coupling transformation of carbon, nitrogen and sulfur in a long-term operated full-scale constructed wetland. Science of The Total Environment 777, 146016. 10.1016/j.scitotenv.2021.146016

Liu, X., Wang, Y., Liu, H., Zhang, Y., Zhou, Q., Wen, X., Guo, W., Zhang, Z., 2024. A systematic review on aquaculture wastewater: Pollutants, impacts, and treatment technology. Environmental Research 262, 119793. 10.1016/j.envres.2024.119793

Louca, S., Parfrey, L.W., Doebeli, M., 2016. Decoupling function and taxonomy in the global ocean microbiome. Science 353, 1272–1277. 10.1126/science.aaf4507

Ma, J., Cui, Y., Li, A., Zou, X., Ma, C., Chen, Z., 2022. Antibiotics and antibiotic resistance genes from wastewater treated in constructed wetlands. Ecological Engineering 177, 106548. 10.1016/j.ecoleng.2022.106548

Ma, X., Song, X., Li, X., Fu, S., Li, M., Liu, Y., 2018. Characterization of Microbial Communities in Pilot-Scale Constructed Wetlands with Salicornia for Treatment of Marine Aquaculture Effluents. Archaea 2018, 7819840. 10.1155/2018/7819840

Martin, M., 2011. Cutadapt removes adapter sequences from high-throughput sequencing reads. EMBnet j. 17, 10. 10.14806/ej.17.1.200

McMurdie, P.J., Holmes, S., 2013. phyloseq: An R Package for Reproducible Interactive Analysis and Graphics of Microbiome Census Data. PLOS ONE 8, e61217. 10.1371/journal.pone.0061217

Melita, M., Amalfitano, S., Preziosi, E., Ghergo, S., Frollini, E., Parrone, D., Zoppini, A., 2022. Redox conditions and a moderate anthropogenic impairment of groundwater quality reflected on the microbial functional traits in a volcanic aquifer. Aquat Sci 85, 3. 10.1007/s00027-022-00899-8

Menzel, P., Ng, K.L., Krogh, A., 2016. Fast and sensitive taxonomic classification for metagenomics with Kaiju. Nat Commun 7, 11257. 10.1038/ncomms11257

Miki, T., Yokokawa, T., Ke, P., Hsieh, I., Hsieh, C., Kume, T., Yoneya, K., Matsui, K., 2018. Statistical recipe for quantifying microbial functional diversity from EcoPlate metabolic profiling. Ecological Research 33, 249–260. 10.1007/s11284-017-1554-0

Moazzem, S., Bhuiyan, M., Muthukumaran, S., Fagan, J., Jegatheesan, V., 2023. Microbiome Wetlands in Nutrient and Contaminant Removal. Curr Pollution Rep 9, 694–709. 10.1007/s40726-023-00280-9

Moschos, S., Kormas, K.Ar., Karayanni, H., 2022. Prokaryotic diversity in marine and freshwater recirculating aquaculture systems. Reviews in Aquaculture 14, 1861–1886. 10.1111/raq.12677

Mustafa, G., Hussain, S., Liu, Y., Ali, I., Liu, J., Bano, H., 2024. Microbiology of wetlands and the carbon cycle in coastal wetland mediated by microorganisms. Science of The Total Environment 954, 175734. 10.1016/j.scitotenv.2024.175734

Niu, S., Li, C., Xie, J., Li, Z., Zhang, K., Wang, G., Xia, Y., Tian, J., Li, H., Xie, W., Gong, W., 2025. Influence of aquaculture practices on microbiota composition and pathogen abundance in pond ecosystems in South China. Water Research X 27, 100302. 10.1016/j.wroa.2025.100302

Nurk, S., Meleshko, D., Korobeynikov, A., Pevzner, P.A., 2017. metaSPAdes: a new versatile metagenomic assembler. Genome Res. 27, 824–834. 10.1101/gr.213959.116

Olson, R.D., Assaf, R., Brettin, T., Conrad, N., Cucinell, C., Davis, J.J., Dempsey, D.M., Dickerman, A., Dietrich, E.M., Kenyon, R.W., Kuscuoglu, M., Lefkowitz, E.J., Lu, J., Machi, D., Macken, C., Mao, C., Niewiadomska, A., Nguyen, M., Olsen, G.J., Overbeek, J.C., Parrello, B., Parrello, V., Porter, J.S., Pusch, G.D., Shukla, M., Singh, I., Stewart, L., Tan, G., Thomas, C., VanOeffelen, M., Vonstein, V., Wallace, Z.S., Warren, A.S., Wattam, A.R., Xia, F., Yoo, H., Zhang, Y., Zmasek, C.M., Scheuermann, R.H., Stevens, R.L., 2023. Introducing the Bacterial and Viral Bioinformatics Resource Center (BV-BRC): a resource combining PATRIC, IRD and ViPR. Nucleic Acids Res 51, D678– D689. 10.1093/nar/gkac1003

Quast, C., Pruesse, E., Yilmaz, P., Gerken, J., Schweer, T., Yarza, P., Peplies, J., Glöckner, F.O., 2013. The SILVA ribosomal RNA gene database project: improved data processing and web-based tools. Nucleic Acids Research 41, D590–D596. 10.1093/nar/gks1219

Quero, G.M., Piredda, R., Basili, M., Maricchiolo, G., Mirto, S., Manini, E., Seyfarth, A.M., Candela, M., Luna, G.M., 2023. Host-associated and Environmental Microbiomes in an Open-Sea Mediterranean Gilthead Sea Bream Fish Farm. Microb Ecol 86, 1319–1330. 10.1007/s00248-022-02120-7

Rana, A., Kobayashi, T., Ralph, T.J., 2021. Planktonic Metabolism and Microbial Carbon Substrate Utilization in Response to Inundation in Semi-Arid Floodplain Wetlands. Journal of Geophysical Research: Biogeosciences 126, e2019JG005571. 10.1029/2019JG005571

Rani, A., Chauhan, M., Kumar Sharma, P., Kumari, M., Mitra, D., Joshi, S., 2024. Microbiological dimensions and functions in constructed wetlands: A review. Current Research in Microbial Sciences 7, 100311. 10.1016/j.crmicr.2024.100311

Sabri, N.A., Schmitt, H., van der Zaan, B.M., Gerritsen, H.W., Rijnaarts, H.H.M., Langenhoff, A.A.M., 2021. Performance of full scale constructed wetlands in removing antibiotics and antibiotic resistance genes. Science of The Total Environment 786, 147368. 10.1016/j.scitotenv.2021.147368

Salomo, S., Münch, C., Röske, I., 2009. Evaluation of the metabolic diversity of microbial communities in four different filter layers of a constructed wetland with vertical flow by Biolog^TM^ analysis. Water Research 43, 4569–4578. 10.1016/j.watres.2009.08.009

Schober, I., Koblitz, J., Sardà Carbasse, J., Ebeling, C., Schmidt, M.L., Podstawka, A., Gupta, R., Ilangovan, V., Chamanara, J., Overmann, J., Reimer, L.C., 2025. BacDive in 2025: the core database for prokaryotic strain data. Nucleic Acids Res 53, D748–D756. 10.1093/nar/gkae959

Shaffer, M., Borton, M.A., Bolduc, B., Faria, J.P., Flynn, R.M., Ghadermazi, P., Edirisinghe, J.N., Wood-Charlson, E.M., Miller, C.S., Chan, S.H.J., Sullivan, M.B., Henry, C.S., Wrighton, K.C., 2023. kb_DRAM: annotation and metabolic profiling of genomes with DRAM in KBase. Bioinformatics 39, btad110. 10.1093/bioinformatics/btad110

Skarżyńska, M., Zając, M., Bomba, A., Bocian, Ł., Kozdruń, W., Polak, M., Wiącek, J., Wasyl, D., 2021. Antimicrobial Resistance Glides in the Sky—Free-Living Birds as a Reservoir of Resistant Escherichia coli With Zoonotic Potential. Front. Microbiol. 12. 10.3389/fmicb.2021.656223

Sleytr, K., Tietz, A., Langergraber, G., Haberl, R., 2007. Investigation of bacterial removal during the filtration process in constructed wetlands. Science of The Total Environment, Contaminants in Natural and Constructed Wetlands: Pollutant Dynamics and Control 380, 173–180. 10.1016/j.scitotenv.2007.03.001

Stefanowicz, A., 2006. The Biolog Plates Technique as a Tool in Ecological Studies of Microbial Communities. Polish Journal of Environmental Studies 15.

Sun, G., Zhu, Y., Saeed, T., Zhang, G., Lu, X., 2012. Nitrogen removal and microbial community profiles in six wetland columns receiving high ammonia load. Chemical Engineering Journal 203, 326–332. 10.1016/j.cej.2012.07.052

Tian, W., Li, Q., Luo, Z., Wu, C., Sun, B., Zhao, D., Chi, S., Cui, Z., Xu, A., Song, Z., 2024. Microbial community structure in a constructed wetland based on a recirculating aquaculture system: Exploring spatio-temporal variations and assembly mechanisms. Marine Environmental Research 197, 106413. 10.1016/j.marenvres.2024.106413

Truu, M., Juhanson, J., Truu, J., 2009. Microbial biomass, activity and community composition in constructed wetlands. Science of The Total Environment, Thematic Papers: Selected papers from the 2007 Wetland Pollutant Dynamics and Control Symposium 407, 3958–3971. 10.1016/j.scitotenv.2008.11.036

Verduzo Garibay, M., Fernández del Castillo, A., de Anda, J., Senés-Guerrero, C., Gradilla-Hernández, M.S., 2022a. Structure and activity of microbial communities in response to environmental, operational, and design factors in constructed wetlands. Int. J. Environ. Sci. Technol. 19, 11587–11612. 10.1007/s13762-021-03719-y

Verduzo Garibay, M., Fernández del Castillo, A., de Anda, J., Senés-Guerrero, C., Gradilla-Hernández, M.S., 2022b. Structure and activity of microbial communities in response to environmental, operational, and design factors in constructed wetlands. Int. J. Environ. Sci. Technol. 19, 11587–11612. 10.1007/s13762-021-03719-y

Vymazal, J., 2022. The Historical Development of Constructed Wetlands for Wastewater Treatment. Land 11, 1–29.

Vymazal, J., Zhao, Y., Mander, Ü., 2021. Recent research challenges in constructed wetlands for wastewater treatment: A review. Ecological Engineering 169, 106318. 10.1016/j.ecoleng.2021.106318

Wang, C., Yu, J., Zhang, J., Zhu, B., Zhao, W., Wang, Z., Yang, T., Yu, C., 2024. A review of factors affecting the soil microbial community structure in wetlands. Environ Sci Pollut Res 31, 46760–46768. 10.1007/s11356-024-34132-w

Wang, J., Long, Y., Yu, G., Wang, G., Zhou, Z., Li, P., Zhang, Y., Yang, K., Wang, S., 2022. A Review on Microorganisms in Constructed Wetlands for Typical Pollutant Removal: Species, Function, and Diversity. Front. Microbiol. 13. 10.3389/fmicb.2022.845725

Wanyan, R., Pan, M., Mai, Z., Xiong, X., Wang, S., Han, Q., Yu, Q., Wang, G., Wu, S., Li, H., 2023. Fate of high-risk antibiotic resistance genes in large-scale aquaculture sediments: Geographical differentiation and corresponding drivers. Science of The Total Environment 905, 167068. 10.1016/j.scitotenv.2023.167068

Weinstein, M.M., Prem, A., Jin, M., Tang, S., Bhasin, J.M., 2019. FIGARO: An efficient and objective tool for optimizing microbiome rRNA gene trimming parameters. 10.1101/610394

Wu, S., Kuschk, P., Wiessner, A., Müller, J., Saad, R.A.B., Dong, R., 2013. Sulphur transformations in constructed wetlands for wastewater treatment: A review. Ecological Engineering 52, 278–289. 10.1016/j.ecoleng.2012.11.003

Wu, Y.-W., Simmons, B.A., Singer, S.W., 2016. MaxBin 2.0: an automated binning algorithm to recover genomes from multiple metagenomic datasets. Bioinformatics 32, 605–607. 10.1093/bioinformatics/btv638

Xiong, R., Li, Y., Gao, X., Li, N., Lou, R., Saeed, L., Huang, J., 2023. Effects of a long-term operation wetland for wastewater treatment on the spatial pattern and function of microbial communities in groundwater. Environmental Research 228, 115929. 10.1016/j.envres.2023.115929

Xu, C., Feng, Y., Li, H., Li, Y., Yao, Y., Wang, J., 2024. Constructed wetlands for mariculture wastewater treatment: From systematic review to improvement measures and insights. Desalination 579, 117505. 10.1016/j.desal.2024.117505

Yang, J., Wang, C., Shu, C., Liu, L., Geng, J., Hu, S., Feng, J., 2013. Marine Sediment Bacteria Harbor Antibiotic Resistance Genes Highly Similar to Those Found in Human Pathogens. Microb Ecol 65, 975–981. 10.1007/s00248-013-0187-2

Yang, R., Fang, J., Cao, Q., Zhao, D., Dong, J., Wang, R., Liu, J., 2021. The content, composition, and influencing factors of organic carbon in the sediments of two types of constructed wetlands. Environ Sci Pollut Res 28, 49206–49219. 10.1007/s11356-021-14134-8

Zhang, H., Chen, R., He, Y., Cao, Z., Zhou, R., Zheng, C., Pan, D., Fang, H., Wu, X., 2025. Integrating livestock and aquatic plant towards mitigating antibiotic resistance transmission from swine wastewater. npj Clean Water 8, 14. 10.1038/s41545-025-00446-6

Zhao, X., Zhang, T., Yang, J., Zhang, H., Yang, L., Li, Q., Hou, N., 2024. Recovery capacity of constructed wetlands in response to multiple disturbances: Microbial interaction perspective. Bioresource Technology 408, 131155. 10.1016/j.biortech.2024.131155

Zhou, S., Liu, B., Zheng, D., Chen, L., Yang, J., 2025. VFDB 2025: an integrated resource for exploring anti-virulence compounds. Nucleic Acids Res 53, D871–D877. 10.1093/nar/gkae968

